# Transcription is ubiquitously terminated in thousands of bidirectional termination zones in yeast

**DOI:** 10.1101/2022.11.02.514956

**Authors:** Gang Zhen, Buki Kwon

## Abstract

Pervasive transcription of eukaryotic genomes requires intricate mechanisms to delineate boundaries for each transcriptional unit. How transcription is efficiently terminated before invading neighboring genes remains an open question. Here, after dissecting the cleavage and polyadenylation landscape using a hybrid approach, we observed thousands of bidirectional termination zones in the genome of *Saccharomyces cerevisiae*. These zones are ∼120 bp wide and terminate transcription from both sense and antisense strands in yeast. They are ubiquitously used as termination sites for both coding and non-coding genes. We suggest that the known transcription termination efficiency element, UAUAUA motifs, serves as the central elements in these zones. Notably, bidirectional termination zones are specifically nucleosome depleted, suggesting chromatin structure plays a key role in the formation of bidirectional termination zones in yeast. Finally, we provide evidence for transcriptional interference in these bidirectional termination zones, and expression level of each cleavage site is influenced by sequence contexts both upstream and downstream. We provide the first global fine-scale picture of transcription termination in a eukaryotic genome.

## Introduction

The yeast genome is pervasively transcribed^1^. Due to the high compactness of the yeast genome, ∼6000 protein-coding genes dispersed in a ∼12 Mb region, there are frequent overlaps between transcripts from adjacent transcriptional units^2^. Therefore, an important question in yeast genomics is how the boundaries of each transcriptional unit are established and maintained in transcription^3^. Transcription initiation has been extensively studied for many years, and mounting evidence suggests that most yeast promoters are bidirectional, generating protein-coding transcripts on the sense strand and cryptic transcripts on the antisense strand^4–6^. However, regulation of transcription termination has been largely overlooked for a long time^7^.

Some genes’ termination sites were known to enable transcription termination from both directions in yeast many years ago^8,9^. Next generation sequencing data revealed that many 3’UTR regions in yeast are shared between convergently transcribed genes^10,11^. Another study found that terminators of tandem genes (genes that align sequentially on the same strand) could also serve as termination sites for cryptic transcription from antisense (AS) strand^12^. These results suggest bidirectional termination is quite common in yeast genome. However, these studies used RNA-seq data, which is not effective at measuring cleavage site positions since most reads do not contain poly(A) tail sequences. In addition, alternative cleavage and polyadenylation (APA) creates transcripts with different termination sites for the same gene. It plays a key role in the diversification of transcriptome^13,14^, and is widely used in the yeast genome^15,16^. How APA affects bidirectional termination is currently unknown. In addition, a recent study found that bidirectional termination of convergently transcribed genes has an influence on 3’UTR length^17^, suggesting it may play an important regulatory role in transcription.

Here, we carried out the first fine-scale genome-wide analysis of transcription termination in yeast. We show that transcription is ubiquitously terminated in bidirectional termination zones in yeast. The long-known efficiency element in yeast 3’UTRs, UAUAUA motif, in fact serves as the bidirectional polyadenylation signal in these termination zones. We found chromatin structure is a key factor in controlling transcription termination in yeast. Finally, we show how these bidirectional termination zones impact expression of individual 3’UTR isoform.

## Results

### A high-resolution map of cleavage and polyadenylation sites in yeast

The yeast transcriptome is extremely complex due to frequent overlaps between transcripts from neighboring genes^2^. In this study, we took a hybrid approach to dissect all cleavage and polyadenylation sites (pAs) by using both 3’-end sequencing data and long read sequencing data (Fig. 1a). First, we employed 3’-seq^18^ to sequence 3’-ends of polyadenylated transcripts, and then mapped all pAs by peak-calling (see Methods). Next, we sought to assign each pA to the right upstream gene. In yeast, there are three major scenarios where a pA could be easily mis-assigned to a wrong gene. First, a pA located within a gene’s coding region (CDR) could be either a truncated transcript of the same gene or is a transcription termination site of the upstream gene (Supplementary Fig. 1a). Second, pAs coming from non-coding RNAs (ncRNAs) could be easily mis-assigned to upstream coding genes (Fig. 1a (plus strand); Supplementary Fig. 1b). Third, poly-cistronic transcripts are frequently found for many pAs (Supplementary Fig. 1c), making proper assignments of these pAs much difficult. To address these challenges, we used long read sequencing data from published datasets^2,19,20^ to help us assign each pA to the right gene (see Methods).

**Figure 1:**
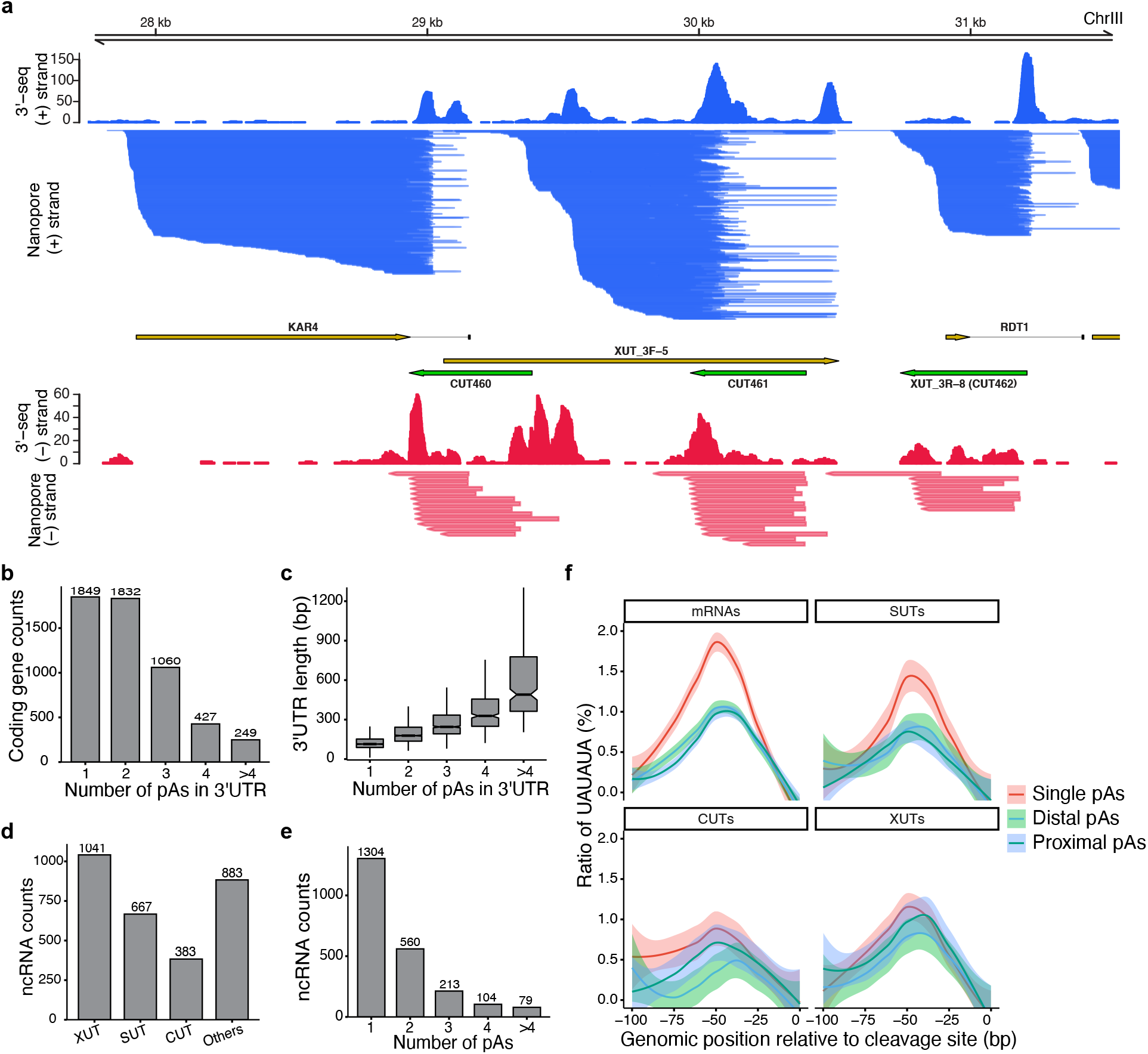
Annotation of cleavage and polyadenylation sites (pAs) in yeast. (a) An illustration of pA annotation by using both 3’-seq and nanopore mRNA-direct sequencing data. (b) Summary of pA annotation results of protein-coding genes. (c) 3’UTR lengths increase with the number of pAs. (d) Annotation of ncRNAs identified in our study. (e) Summary of pA annotation results for ncRNAs. (f) UAUAUA motif distribution among single, distal, and proximal pAs in mRNAs and ncRNAs.

Finally, 13,720 high-qualify pAs were assigned to 5,417 protein-coding gene. Among these pAs, 11,762 (85.7%) fall into 3’UTRs, while the rest are located in CDRs (Supplementary Fig. 1e). Due to poly-cistronic reads, 634 pAs (578 in 3’UTRs and 56 in CDRs) could be assigned to more than one protein-coding genes. At the gene level, 3,568 protein-coding genes (65.9%) are associated with more than one pA (Fig. 1b), confirming widespread APA in yeast^15,16^. Importantly, since we included only those pAs that were also supported by long reads, the real number of genes with multiple pAs may be even larger as long read sequencing doesn’t provide the same read depth as standard RNA-seq. We determined that the median length of 3’UTRs in yeast is 179 bp (average length: 247.6 bp; Supplementary Fig. 1f), which is slightly shorter than that of *Schizosaccharomyces pombe* (median length: 187.5 bp)^21^ and is much longer than the multi-cellular organism *Caenorhabditis elegans* (median length: 140 bp)^22^. In yeast, the length of 3’UTR strongly correlates with the number of pAs (Fig. 1c), suggesting a relationship between 3’UTR length and transcriptome complexity.

Pervasive transcription in yeast genome generates thousands of ncRNAs, many of which have been shown to play important regulatory or functional roles^23–25^. In contrast to coding genes, APA in ncRNAs has never been systematically characterized in yeast. The genomic origins of ncRNAs are highly heterogenous^26,27^, requiring the identification of genomic boundaries for each ncRNA a prerequisite prior to APA analysis. We first used those un-assigned pAs from previous analysis as seeds, and then used long reads to re-construct the full-length of ncRNAs (see Methods). Thus, we obtained exact genomic locations for 2,260 high-confidence ncRNAs. By mapping to published ncRNA datasets in yeast^4,28^, 1,377 of these ncRNAs were assigned to at least one of the three pre-established major classes of ncRNAs (SUTs/CUTs/XUTs; Fig. 1d).

Annotation result shows that APA is also widespread in ncRNAs. For example, *SRG1*, which is generated from the 5’UTR of the *SER3* gene and negatively regulates expression of *SER3* under serine-rich condition in yeast^29^, has three validated pAs (Supplementary Fig. 2a). *IRT1*, which can inhibit expression of the downstream gene *IME1* and thus play a role in the regulation of meiosis in yeast^30^, has four pAs (Supplementary Fig. 2b). In addition to these stable transcripts (SUTs), we also observed APA in the other two classes of unstable ncRNAs, XUTs and CUTs (Supplementary Figs. 2c-f). In total, 956 of the 2260 ncRNAs (42.3%) have at least two pAs (Fig. 1e). Among different classes of ncRNAs, SUTs have the highest multi-pA ratio (59.4%), while CUTs have the lowest ratio (34.3%) (Supplementary Fig. 2h). The decreasing multi-pA ratios from SUTs to CUTs correlate well with their decreasing expression levels (Supplementary Fig. 2i), indicating the real multi-pA ratios of XUTs and CUTs may be even higher since some of their transcripts may be quickly degraded before being detected by sequencing.

In yeast, the canonical polyadenylation signal (PAS) AAUAAA, which is commonly found in vertebrates, is highly degenerate, while a UAUAUA motif, known as efficiency element, is strongly enriched in a region 30-100 nt upstream of cleavage site^16,31^. The UAUAUA motif has been known to play a key role in cleavage and polyadenylation^32,33^, and it has also been shown to affect mRNA abundance^34,35^. In our dataset, in addition to its effect on expression levels (Supplementary Fig. 3a), we found the location of UAUAUA motif could affect cleavage strength (see Methods), of which pAs tend to have high cleavage strengths when UAUAUA motifs are located ∼30bp upstream of major cleavage site (CS refers to this major site hereafter, Supplementary Fig. 3c). This result suggests that relative position between UAUAUA motif and CS is a key factor in controlling 3’-end processing efficiency in yeast. Next, we tried to find out whether there is any difference in the distribution of UAUAUA motifs among pAs of single- and multi-UTR genes, and among different kinds of transcripts. In mRNAs, we found that pAs of single-UTR genes are more likely to be preceded by UAUAUA motifs than those of multi-UTR genes (Fig. 1f, Supplementary Fig. 3e). In comparison, we did not observe significant differences in distribution of canonical AAUAAA motifs between single- and multi-UTR genes in mRNAs (Supplementary Figs. 3d,e). These results indicate that high ratio of UAUAUA motifs may help single-UTR genes achieve precise usage of one pA. Among different classes of ncRNAs, SUTs have similar differences in the distribution of UAUAUA motifs between single- and multi-pA genes as mRNAs, which is in concordance with the fact that mRNAs and SUTs share similar cleavage and polyadenylation mechanism^36^, while no such difference was observed in the other two classes of ncRNAs (Fig. 1f, Supplementary Fig. 3e).

### Transcription is ubiquitously terminated in bidirectional termination zones in yeast

Many convergently transcribed protein-coding genes were found to share their 3’UTR regions^10,11^. In our new dataset of pAs in yeast, among the 1396 high-quality convergent gene pairs (see Methods), 1275 of them (91.3%) were found to share their 3’UTRs. This number was much greater than those previously reported (275^10^ and 645^11^ pairs). This result indicates that bidirectional termination is a general phenomenon for convergently transcribed genes in yeast. Interestingly, for many of these gene pairs, their pAs seems also to be paired (Supplementary Figs. 4a,b). Therefore, we raised a hypothesis that, for each convergent gene pair, positions of CSs on one strand are correlated to the positions of CSs on the other strand. To test it, we visualized the 3’-end distribution of AS reads relative to the CSs on the sense strand. Considering antisense artifacts are common in strand-specific RNA-seq protocols^37^, we used a long read dataset^19^ generated by nanopore mRNA-direct sequencing technology for this analysis. In order to determine how APA affects bidirectional termination, we carried out this analysis for single-UTR and multi-UTR genes separately. For single-UTR genes, interestingly, regardless of 3’UTR length, the 3’-ends of AS reads are strongly enriched in a region 60-120 bp upstream of CSs, suggesting that termination of AS transcription is strictly confined by sense CSs (Fig. 2a). This result also reveals existence of these ∼120 bp wide bidirectional termination zones in yeast genome. For multi-UTR genes, when aligned by distal CSs, 3’-ends of AS reads are also enriched shortly upstream (Fig. 2b). However, unlike single-UTR genes, the distribution of AS 3’-ends for multi-UTR genes expands further upstream of distal CSs and strongly correlates with the positions of proximal CSs on sense strand (Fig. 2b). This result indicates that, for multi-UTR genes, the distribution of AS CSs is confined by both the proximal and distal CSs on the sense strand. It also indicates that both proximal and distal CSs are located within bidirectional termination zones. To confirm these findings, we repeated this analysis using three other RNA-seq datasets generated by different technologies (3’READS^38^, Tif-seq^2^, and Pab1 (a poly(A)-binding protein involved in cleavage and polyadenylation in yeast^39,40^) CLIP-seq^36^). The 3’-end distributions of AS reads in all three datasets follow exactly the same patterns as described above (Supplementary Figs. 4c-h), demonstrating that these bidirectional termination zones are highly reproducible in yeast genome.

**Figure 2:**
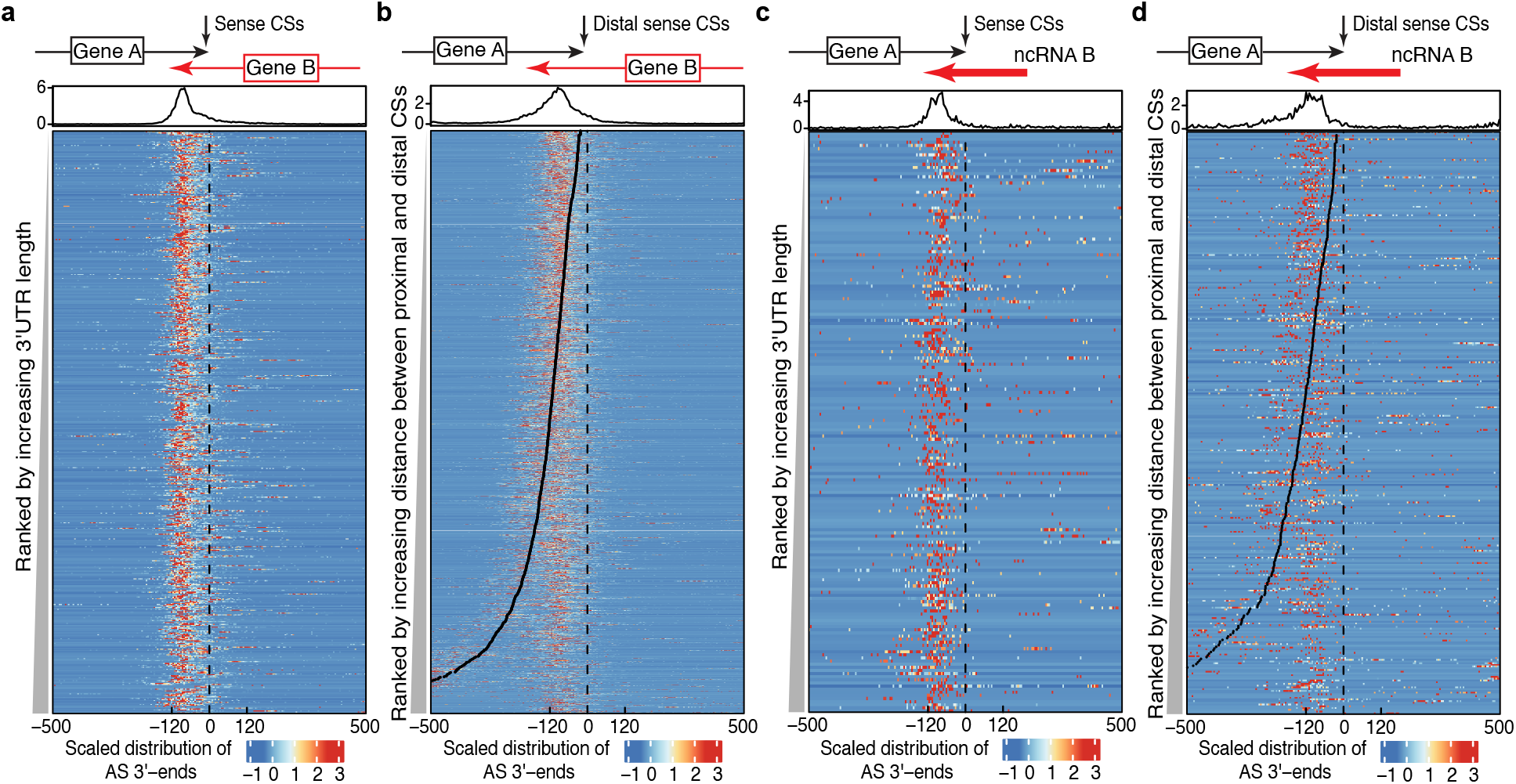
Genome-wide distribution of bidirectional termination zones in yeast. (a, c) Heatmaps show 3’-end distribution of AS nanopore mRNA-direct sequencing reads in a 1000 bp window centered at CSs (black dashed lines) of convergent (a) and tandem (c) single-UTR genes with bidirectional terminators. (b, d) Heatmaps show 3’-end distribution of AS nanopore reads centered at distal CSs (black dashed lines) of convergent (b) and tandem (d) multi-UTR genes with bidirectional terminators. Black dots in b & d represent positions of proximal CSs.

For the remaining 121 convergent gene pairs that don’t share their 3’UTR regions, we noticed that some of these genes share their 3’UTRs with termination sites of AS ncRNAs (Supplementary Fig. 5a). This finding suggests that 3’UTRs of these genes may also contain bidirectional termination zones. After checking our atlas of ncRNAs, among these 242 convergent genes, 81 of them have an AS ncRNA downstream, and 70 of them share their 3’UTRs with pAs of AS ncRNAs. Even though the expression levels of ncRNAs are much lower than mRNAs (Supplementary Fig. 2i), we could still see bidirectional termination patterns for these 70 genes (Supplementary Figs. 5b,c). For the other 172 genes, we confirmed that the termination sites of these genes are indeed unidirectional by visualizing the 3’-end distributions of AS reads across all four RNA-seq datasets (Supplementary Fig. 6).

For tandem genes, it was reported that they could share their 3’UTRs with terminators of AS ncRNAs^12^. In our analysis, among 2,827 tandem genes, 843 of them have an AS ncRNA downstream, and 704 of them share their termination regions with AS ncRNAs (see examples in Supplementary Figs. 7a,b). For single-UTR tandem genes, regardless of their 3’UTR length, AS 3’-ends are strongly enriched in a region about 60-120 bp upstream of CSs (Fig. 2c). For multi-UTR tandem genes, the distributions of AS 3’-ends are strongly correlated with both positions of proximal and distal CSs on sense strand (Figs. 2d). These AS 3’-end distribution patterns of tandem genes are highly consistent with those observed in convergent genes, suggesting that bidirectional termination zones are ubiquitous in the yeast genome. Interestingly, for the remaining 2,123 tandem genes that do not share their termination zones with any validated AS ncRNAs, AS 3’-ends are also enriched upstream of sense CSs, even though the signals are much weaker (Supplementary Fig. 8). This result suggests that some of these tandem genes may also have bidirectional termination zones, which are probably shared with some highly unstable ncRNAs generated by pervasive transcription.

### UAUAUA motifs are the central elements in bidirectional termination zones

In transcription, polyadenylation signals (PASs) are required for mRNA 3’-end formation in eukaryotes^41^. With regards to bidirectional termination zones, an interesting question is what type of PAS enables a bidirectional termination pattern. We focused on the UAUAUA motif (and its two important variants: UACAUA and UAUGUA) for three main reasons: 1) these motifs play key roles in cleavage and polyadenylation^32,33^ and could affect genes’ expression levels^34,35^ in yeast; 2) these motifs are highly enriched in bidirectional termination zones (Fig. 1f); 3) these motifs are palindromic, which suggests they could be used for both sense and AS transcription termination. Computation modeling analysis suggested that UAUAUA motif could terminate transcription from both directions^3^. In bidirectional termination zones, when aligned by the position of UAUAUA motifs, across all 3 groups of pAs (single/distal/proximal), the distributions of sense and AS 3’-ends are highly symmetrical (Fig. 3a, Supplementary Fig. 9a), confirming that UAUAUA motifs serve as bidirectional PASs^3^. Our previous analysis shows that UAUAUA motifs influence cleavage strengths (Supplementary Fig. 3c). Interestingly, we found that when there is no UAUAUA motif in bidirectional termination zones, both cleavage sites on sense and AS strands have decreased cleavage strengths (Fig. 3b), suggesting that the UAUAUA motif could affect cleavage strengths on both strands simultaneously. At genome level, for convergent genes, if their 3’UTRs are overlapping, there are two adjacent 3’UTRs on opposite strands. Since 3’UTRs have very special nucleotide composition (highly AU enriched), we expect the nucleotide composition on both sides of the UAUAUA motif to be reverse complemented, which is indeed the case (Fig. 3c, Supplementary Fig. 9b). These results suggest that UAUAUA motifs are the central elements in bidirectional termination zones.

**Figure 3:**
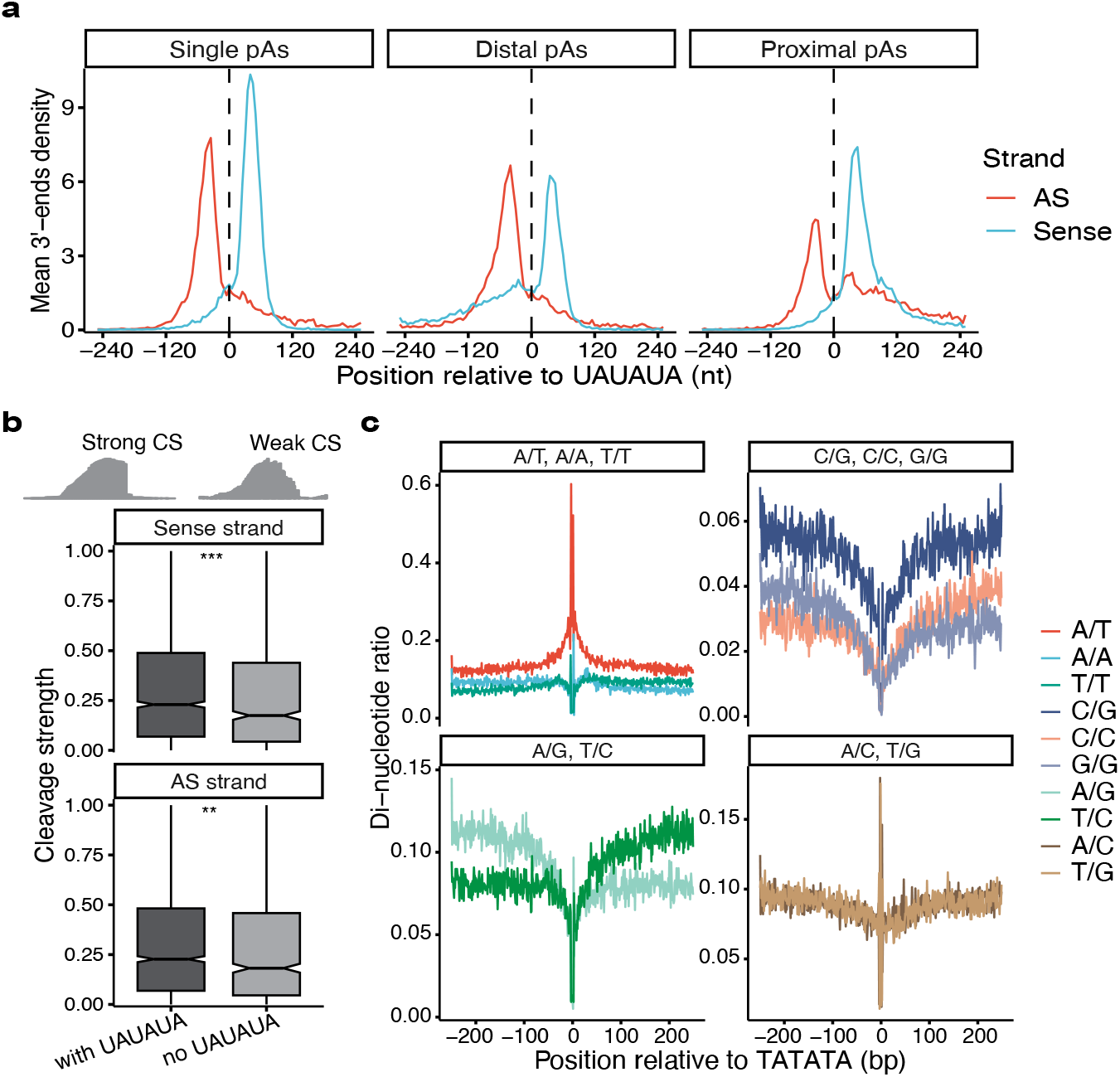
UAUAUA motifs are the central elements in bidirectional termination zones. (a) 3’-ends of sense and AS nanopore mRNA-direct sequencing reads are symmetrically distributed on both sides of UAUAUA motifs in bidirectional termination zones of convergent genes. (b) existence of UAUAUA motif increases cleavage strength of both sense and AS pAs. (c) di-nucleotide compositions on both sides of UAUAUA motifs are reverse complemented.

### Bidirectional termination zones are specifically nucleosome-depleted

Yeast 3’UTRs are known to be nucleosome depleted^42–44^, and it was suggested to play a certain role in 3’-end formation^45^. With the help of our high-resolution map of pAs in yeast, we wanted to figure out whether there is a relationship between nucleosome depleted regions (NDRs) and bidirectional termination zones. We therefore analyzed a high-quality nucleosome occupancy profile generated by an H3 chemical cleavage mapping method^46^, which can eliminate sequence bias commonly found in MNase mapping method^47^. For single-UTR convergent overlapping genes, we found that NDRs start at about 120 bp upstream and end shortly downstream of CSs (Fig. 4a). Interestingly, these NDRs match almost perfectly well with bidirectional termination zones of single-UTR genes (Fig. 2a). For multi-UTR convergent overlapping genes, NDRs expand with the increasing distance between proximal and distal CSs (Fig. 4b), whose pattern also matches perfectly well with bidirectional termination zones (Fig. 2b). However, for the 172 unidirectional terminators (Supplementary Figs. 6), NDRs are not confined by cleavage sites and extend further downstream for both single- and multi-UTR genes (Supplementary Figs. 10a,b). In summary, bidirectional termination zones of convergent overlapping genes are demarcated by NDRs, suggesting a key role of chromatin structure in the formation of these bidirectional termination zones. This result also suggests transcription termination is strongly dictated by chromatin structure in yeast.

**Figure 4:**
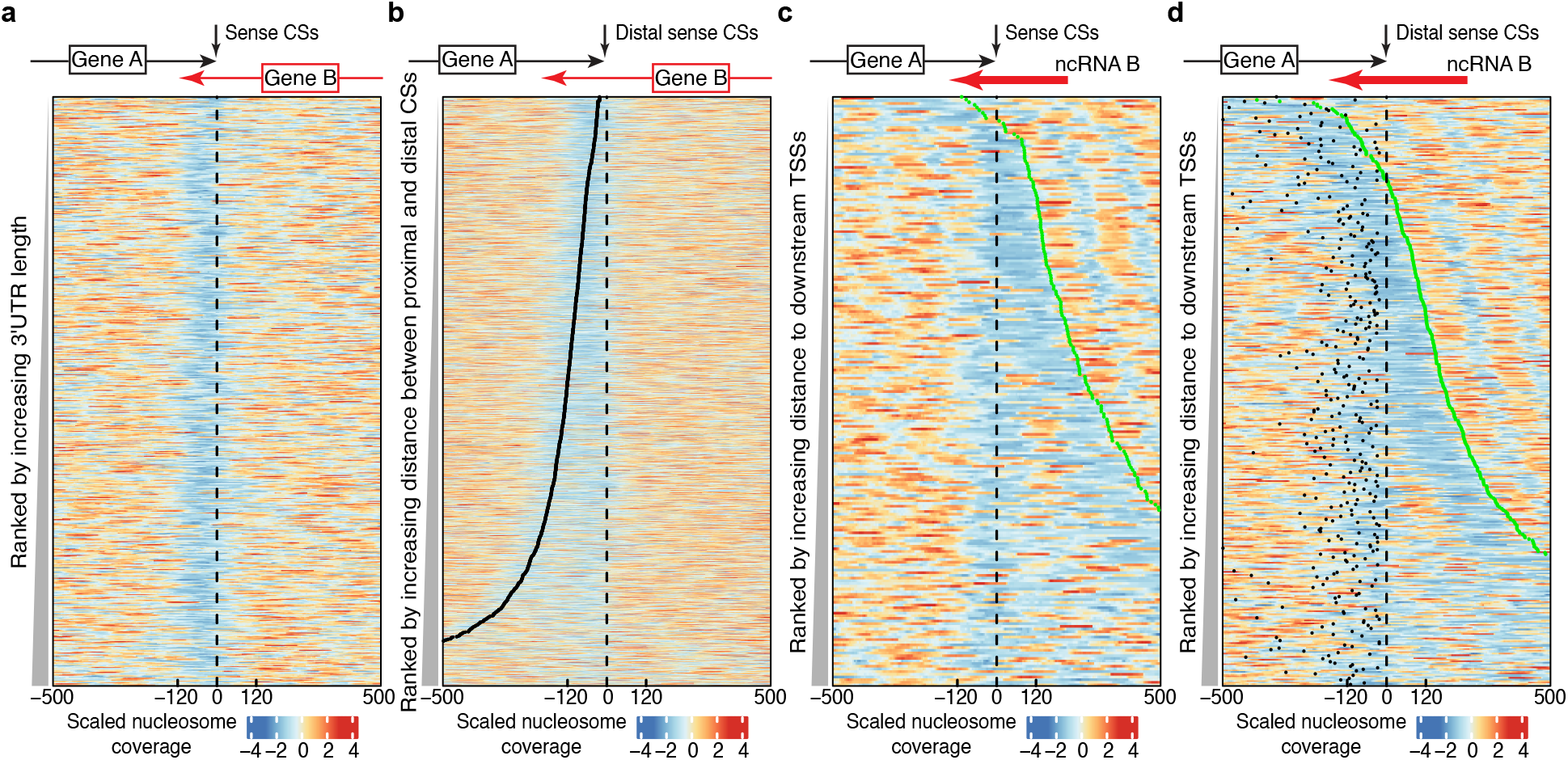
Bidirectional termination zones are specifically nucleosome depleted. (a, c) Heatmaps show nucleosome occupancy in a 1000 bp window centered at CSs of convergent (a) and tandem (c) single-UTR genes with bidirectional terminators. (b, d) Heatmaps show nucleosome occupancy in a 1000 bp window centered at distal CSs of convergent (b) and tandem (d) multi-UTR genes with bidirectional terminators. Black dots in b & d represents positions of proximal CSs. Green dots in c & d represents positions of downstream TSSs. Coordinates of TSSs are downloaded from published paper^48^.

For tandem genes, due to the compactness of yeast genome, we expect that there will be a downstream transcription start site (TSS) nearby for each termination site. As is well known, TSSs are located in regions that are highly nucleosome depleted^43,44^. Therefore, for terminators of tandem genes, an interesting question is whether they share the same NDRs with downstream TSSs. We found that, for single-UTR tandem genes with bidirectional termination zones, their CSs rarely share NDRs with downstream TSSs (Fig. 4c). We observed that 50 of 221 (22.6%) such CSs were in the same NDRs as downstream TSSs (0-150 bp distance). In comparison, 281 of 785 unidirectional single-UTR tandem genes (35.8%) have their CSs in the same NDRs as downstream TSSs (Supplementary Fig. 10c). For multi-UTR tandem genes, both proximal and distal CSs of bidirectional terminators tend to be less likely to share NDRs with downstream TSSs than those of unidirectional terminators (proximal: 13.8% vs 20.5%; distal: 28.8% vs 40.2%; Fig. 4d, Supplementary Fig. 10d). In general, for both single- and multi-UTR tandem genes, CSs of bidirectional terminators are located much further away from downstream TSSs than those of unidirectional terminators (Supplementary Fig. 10e). These results suggest that, for tandem pAs, bidirectional termination zones require specifically formed NDRs. They also suggest that distance between transcriptional unit is an important factor in controlling pervasive transcription.

### Transcriptional interference region hidden in bidirectional termination zones

Since bidirectional termination zones are ubiquitously distributed in the yeast genome (Fig. 2), we want to know whether they could affect expression of two genes sharing a termination zone. Due to the low expression of ncRNAs (Supplementary Fig. 2i), we focused on the 1,275 convergent overlapping coding gene pairs in this analysis. By calculating Spearman’s correlation coefficient between two convergent genes’ expression levels, we found only one significant yet very weak correlation (Spearman’s rho = -0.078, p = 0.005) in the two recent published 3’-READs datasets (Supplementary Fig. 11a). To confirm this result, we further analyzed two strand-specific RNA-seq datasets^49,50^. Correlation analysis showed that there is no expression correlation between convergently transcribed genes in these two datasets (Supplementary Fig. 11b). These results indicate that the bidirectional termination zones in yeast have either little or no influence on expression at the gene level.

Next, we wanted to know whether the influence on expression in fact happens between sense and AS pAs in the same bidirectional termination zones. We grouped all pairs of sense and AS pAs based on the distance between their cleavage sites. Within each group, we calculated the Spearman’s rho between sense and AS pAs’ expression levels. To increase the robustness of the result, we repeated this analysis in five public datasets (three kinds of technologies). Interestingly, we found that there is a strong negative correlation between sense and AS pAs’ expression levels, yet this correlation is highly dependent on the distance between their CSs (Fig. 5a). When two sense and AS pAs have either no or very short overlap (position > -20), there is very weak correlation between their expression levels. However, when the AS CS is located about 20-60 bp upstream of sense CS, a strong negative correlation is formed between their expression levels. When the AS CS is located > 60 bp upstream of sense CS, the negative correlation quickly disappears. This correlation pattern is consistent across three kinds of pAs (single/distal/proximal). This result suggests there is a transcriptional interference region hidden in bidirectional termination zones.

**Figure 5:**
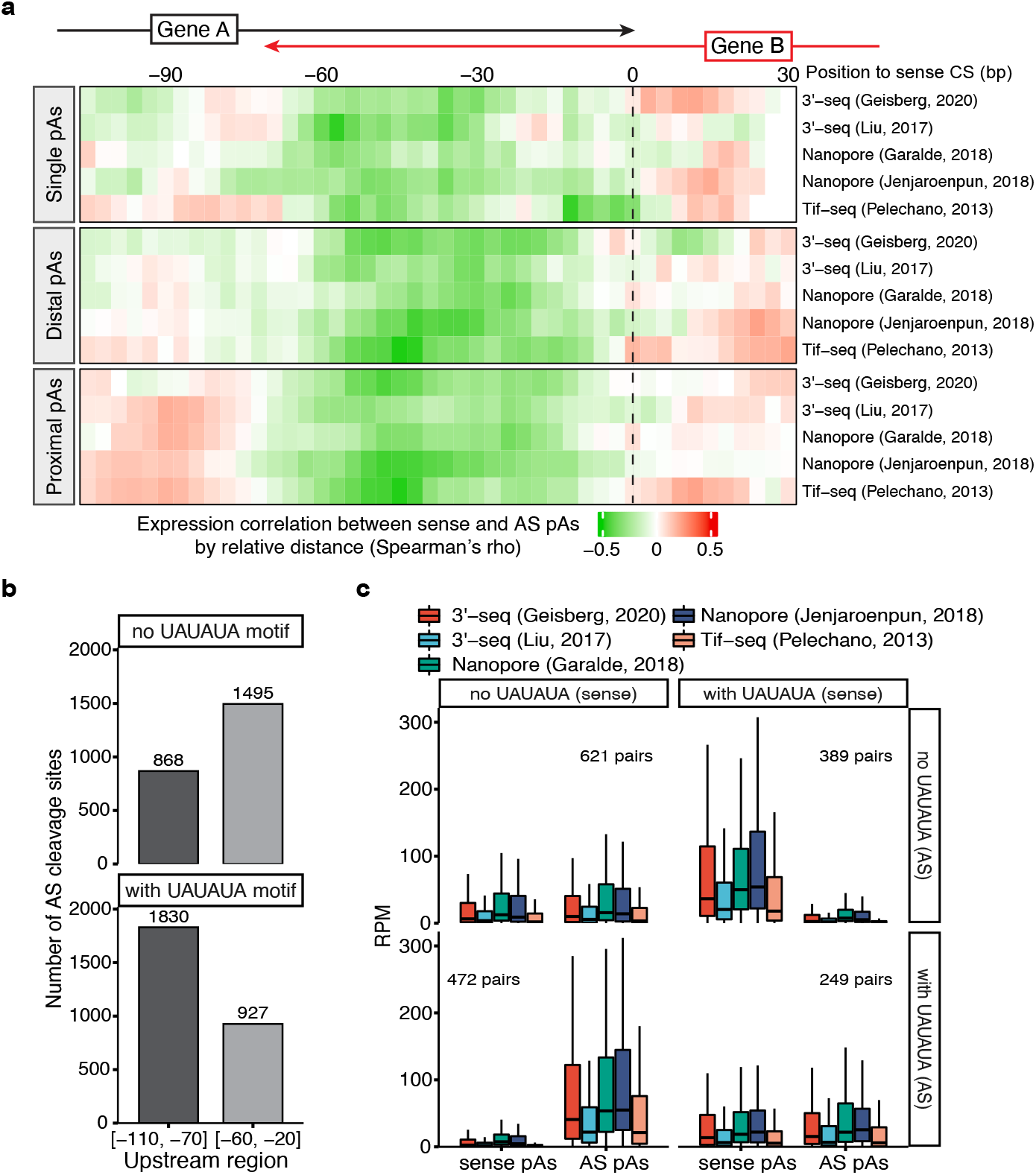
Transcriptional interference hidden in bidirectional termination zones. (a) there is a transcriptional interference region ranging from 20 to 60 bp upstream of sense CSs. Black dashed line represents the positions of sense CSs. Labels above heatmap represent AS CSs’ positions relative to sense CSs. (b) UAUAUA motifs in the transcriptional interference region have an impact in the distribution of AS CSs. (c) when AS CSs fall within transcriptional interference region, expression levels of both sense and AS pAs were greatly influenced by UAUAUA motifs in both upstream and downstream regions of CSs. RPM: reads per million.

### Expression levels of 3’UTR isoforms are influenced by sequence contexts both upstream and downstream of cleavage sites

Interestingly, this transcriptional interference region is also the place where UAUAUA motif is highly enriched in (Fig. 1f). To find out whether the UAUAUA motif plays a role in the transcriptional interference between sense and AS pAs, we compared the distributions of AS CSs between sense pAs with and without UAUAUA motifs. For sense pAs that don’t have UAUAUA motifs, more AS CSs fall into [-60, -20] region compared to those that fall into [-110, -70] region (Fig. 5b, up panel). However, if sense pAs contain UAUAUA motifs, most AS CSs will fall into [-110, -70] region (Fig. 5b, bottom panel). These results suggest that UAUAUA motifs can influence the distribution of AS CSs and help them to avoid the interference zone.

We noticed that, for the 1,495 AS CSs that fall into [-60, -20] region of sense pAs without UAUAUA motifs, 43% of them contain UAUAUA motifs, most of which are located downstream of sense CSs (Supplementary Fig. 11d). This result indicates that, for those pAs in bidirectional termination zones, both genomic sequences upstream and downstream of CSs may affect expression levels due to their bidirectional nature. To investigate the relationship between transcript isoform expression and the presence of the UAUAUA motif, we examined 1731 pA pairs of which AS CSs fall into [-60, -20] region of sense CSs, and separated them into four groups based on presence of UAUAUA motifs for both sense and AS pAs. Then we compared the expression levels between sense and AS pAs in each group (Fig. 5c). When both sense and AS pAs either have or do not have UAUAUA motifs, we did not observe great differences in expression levels between them in any of the five datasets we analyzed. However, when only one of the overlapping sense and AS pAs has a UAUAUA motif, we observed a very strong discrepancy in expression levels between them. In this case, it’s those pAs with UAUAUA motifs that have the highest expression levels among all groups, while those on the opposite strand and without UAUAUA motifs have the lowest. These findings suggest that, for pAs in bidirectional termination zones, both sequence contexts upstream and downstream of the CSs could have a strong impact on their expression levels.

## Discussion

Yeast transcriptome is highly heterogeneous^2^, and antisense transcription is very common^4,16^. In this study, by using both 3’-end sequencing and long read sequencing data, we generated a high-quality single-nucleotide resolution map of cleavage and polyadenylation events in yeast.

This approach produced 13,720 pAs, revealing widespread APA both in coding and non-coding genes. This result suggests that yeast transcriptome is much more diversified than previously thought. By digging into this map of termination sites in yeast, we found that overlapping of 3’UTRs between convergent genes is much more common than previous studies (> 90% of convergent genes). It suggests that sharing of termination sites between neighboring genes is an important way to deal with the limited space in yeast genome. Our further analysis revealed a surprising picture in yeast transcriptome: there are thousands of bidirectional termination zones, which are about 120 bp wide, that can terminate transcription from both directions and effectively prevent transcription invasion of neighboring genes. These zones are used as termination sites not only for coding genes, but also for non-coding genes, suggesting they are used ubiquitously for transcription termination in yeast.

There have been two mysteries in yeast 3’UTRs for a long time. The first one is a highly enrichment of UAUAUA motifs shortly upstream of cleavage sites^16,31^, and the second one is nucleosome depletion^42–44^. Our analysis provides strong evidence suggesting that these UAUAUA motifs are the central elements in the bidirectional termination zones, and they can affect expression termination in both directions (Fig. 3). This is an interesting finding as yeast genome has evolved such an efficient way to save genomic space by using only one PAS instead of two. Then, after revisiting nucleosome coverage in yeast 3’UTRs, we found that the nucleosome depleted regions correspond perfectly well with the bidirectional termination zones (Fig. 4). This result strongly indicates that chromatin structure plays a key role in transcription termination in yeast. Considering bidirectional termination zones are specifically nucleosome depleted, they might be a group of special chromatin structures in yeast genome. Interestingly, previous analysis showed that cohesin is highly enriched in intergenic regions between convergent genes in yeast^51,52^, suggesting that bidirectional termination zones may be part of higher-order genome structures (like chromatin loops^53^). How these chromatin structures are formed and how they influence transcription termination are questions that remain to be answered.

Yeast 3’UTRs have been proven to be able to influence gene expression, especially for regions that are UA enriched^34,35^. In this study, we found that there is a transcriptional interference region ranging from 20 to 60 bp upstream of CSs in bidirectional termination zones (Fig. 5a). This transcriptional interference happens at 3’UTR isoform level, not at gene level, suggesting there is transcriptional regulation in yeast 3’UTRs hidden from regular expression analysis at gene level. Interestingly, this transcriptional interference region coincides with the region where UAUAUA motif is highly enriched. Considering UAUAUA motifs have been shown to influence gene expression in yeast^35^, we speculated that they should play a role in the formation of the transcriptional interference region. Indeed, when AS CSs fall into the transcriptional interference region of sense CSs, and when only one of them harbors UAUAUA motif, there is a great discrepancy between expression levels of sense and AS pAs (Fig. 5c). In contrast, when neither or both the sense and AS CSs have UAUAUA motifs, there is not obvious difference between their expression levels. These findings suggest that the sequence contexts both upstream and downstream of cleavage sites in bidirectional termination zones may disrupt the expression level balance between sense and AS pAs.

In this study, we revealed how transcription is effectively terminated by bidirectional termination zones in a highly compact genome. There are plenty of other genomes that also have high gene density (especially for prokaryotes)^54^, and bidirectional terminators have also been reported in bacteria^55^. Therefore, we speculate such bidirectional termination zones may also exist in other compact genomes and serve as key structures to cope with limited genomic space. Future studies on bidirectional termination zones will help us to better understand how transcription is regulated on a genome-wide scale.

## Methods

### Yeast RNA samples

*Saccharomyces cerevisiae* cultures were treated in different stress conditions (diauxic shift, DNA damage, and heatshock) and were sampled four times during metabolic cycle. A detailed description of this yeast 3’-seq dataset will be provided in a separate manuscript.

### Processing of 3’-seq data

Raw reads from each experiment were first processed by trimmomatic (version: 0.39)^56^ to trim low-quality ends (average quality per base < 15, 4 bp window), and 3’-adapters were removed by cutadapt (version: 2.3)^57^. To ensure identification of real cleavage and polyadenylation events (not internal priming), we refer to the analysis pipeline used by Geisberg et al., 2020^38^. First, we selected those reads with at least two terminal adenosines. We counted the number of terminal adenosines for each read, appending this number to the read ID for future reference. Then all terminal adenosines were trimmed off from each read, and remaining sequence was mapped to yeast genome (version: sacCer3) using hisat2 (version: 2.1.0)^58^. Only reads with high mapping quality (mapq > 30) were kept for downstream analysis. We counted the number of consecutive adenosines in genomic sequence immediately downstream of each mapped read, keeping only those reads whose terminal adenosines (labeled in their IDs) exceeds the number of consecutive adenosines in adjacent genomic regions. Ultimately, we obtained 34.9 million high-confident polyadenylated reads out of all 80.1 million reads.

### Identification of polyadenylation sites

All high-confidence polyadenylated reads from previous step were pooled together for peak calling using CLIPanalyze (version: 0.0.10) as described by Lianoglou et al., 2013^18^. Theoretically, each peak corresponds to one cleavage and polyadenylation site (pA). Due to the “wiggle” nature of pA^59^, within one peak, there is a major site where most read 3’-ends fall into, while there are also some minor sites. We used a 3-bp sliding window to find the major site (where most 3’-ends fall into) within each peak, defining the middle base of this major site as the coordinate of cleavage site for each pA. In yeast genome, sometimes two pAs can be closely spaced to each other, enabling only one peak to be called instead of two. To separate these closely spaced pAs, we propose that if we remove reads belonging to the major pA, the signal of second pA could be detected using the same peak calling process. Therefore, we removed all reads whose 3’-ends fall into a 31-bp window centered at the major cleavage site for each peak. For the remaining reads, we did the second round of peak calling and identification of cleavage sites. For this new set of cleavage sites, only those that are at least 20 bp away from any other cleavage sites were kept. Finally, among the 47092 peaks found in the first round of peak calling, 5134 of them were split into 10807 pAs.

### Assigning each pA to upstream gene

Due to frequent overlaps among adjacent transcripts in yeast, it is not straightforward to assign each pA to the correct upstream gene (see first part of Results for the three scenarios where one pA could be easily mis-assigned). We downloaded and processed three published long-read sequencing datasets (two nanopore direct mRNA sequencing dataset^19,20^, and one Tif-Seq dataset^2^) to help us assign each pA to the right upstream gene. In the first step, we assigned promoter regions of five upstream genes to each pA (we define promoter region as starting from 1 bp upstream of start codon and ending after invading 10% of upstream gene’s pA (same strand), with a maximum length of 2 kb). Second step, we counted how many long reads could connect the pA to each of the five upstream promoters. Third step, each pA was assigned to the closest promoter that was supported by at least two long reads. Fourth step, for those pAs that reside in coding regions, to tell whether they generate truncated transcripts or are in fact termination sites of upstream genes, we compared the number of long reads that connect to the two promoters. If the number of long reads connecting to the upstream promoter is at least two times the number of long reads connecting to the promoter of the very gene in which the pA resides, this pA will be reassigned to the upstream gene. Finally, if one pA could be assigned to at least two promoters (each was supported by at least 2 long reads), it indicates that this pA was contributed by polycistronic transcripts.

### Mis-annotated TSSs

Previous analysis found that 150 transcription start sites (TSSs) are located downstream of the annotated start codon (for example: CBP4) in yeast^48^. For these genes, we define the start positions of their promoters as 15 bp downstream of the mapped TSSs.

### Dubious and overlapping genes

Each pA will be first assigned to unambiguous genes. Only those un-assigned pAs will be assigned to dubious genes in the second round of assignment. For two genes overlapping each other on the same strand, even with the help of long reads, it is difficult to accurately assign the right pAs to each of them. Therefore, pAs downstream of these overlapping genes were filtered out from further analysis.

### Reconstruction of non-coding RNAs

First step, for those pAs that could not be assigned to any upstream protein-coding genes, we kept those that have at least five long reads whose 3’-ends mapped to each pA as seeds. Second step, full-length transcript was constructed using long reads. Third step, those overlapping transcripts on the same strand were merged into one non-coding RNA. Final step, since the transcriptome of yeast is highly heterogenous and it is extremely hard to separate overlapping transcripts from adjacent genes, we manually check these ncRNAs by visualizing nanopore mRNA direct sequencing data to ensure the overall quality is good.

### Processing of other data

#### Nanopore mRNA direct sequencing data

Raw reads were mapped to yeast genome (version: sacCer3) using minimap2 (version: 2.17-r941; preset: -x splice; k-mer size: 14)^60^. Mapped reads were filtered by mapping quality: mapq > 45. Filtered bam files were converted to bed format using bedtools (bamtobed)^61^ for downstream analysis.

#### Tif-Seq data

In pA assignment step, to minimize the influence of PCR duplicates in Tif-Seq data, for those reads with identical 5’- and 3’-end coordinates, only one was kept. In pA expression quantification step, all reads were used.

#### Strand-specific RNA-seq

Raw reads from each dataset were first processed by trimmomatic (version: 0.39) to trim low-quality ends (average quality per base < 15, 4 bp window). Filtered reads were mapped to yeast genome (version: sacCer3) using hisat2 (version: 2.1.0). Reads mapped to each gene (CDR + shortest 3’UTR isoform) were counted by featureCounts (version: 1.6.4)^62^.

#### 3’READS data

First four random bases in each raw read were trimmed by Perl script. Low-quality ends were trimmed by trimmomatic (version: 0.39). Poly(T) tails were trimmed by cutadapt (version: 2.3). After these steps, first 20 bases in each read were extracted and mapped to yeast genome (version: sacCer3) by hisat2 (version: 2.1.0). Raw reads generated by 3’READs technique correspond to antisense strand of transcripts^63^. Therefore, after mapping to yeast genome, strand information for each read was shifted to the opposite strand.

#### Pab1 CLIP-seq

Low-quality ends of raw reads were trimmed by trimmomatic (version: 0.39). First 3 random bases were trimmed using a custom Perl script. Barcodes for multiplexing at 5’ ends and adapters at 3’ ends were both trimmed by cutadapt (version: 2.3). Processed reads were mapped to yeast genome (version: sacCer3) by hisat2 (version: 2.1.0).

#### Nucleosome coverage data

Bigwig files containing nucleosome coverage information were downloaded from GEO^46^ and converted to bedGraph format using UCSC utility tool bigWigToBedGraph before downstream analysis.

#### Calculation of cleavage strength

Cleavage and polyadenylation happens in a ’wiggle’ pattern^59^, which means a typical pA usually consists of several cleavage sites (CSs), of which most transcripts are cleaved at a major site. Here we define cleavage strength as the ratio of reads whose 3’-ends map to the major CS compared to those mapping to the whole pA. For each pA, we used a 3-bp sliding window (1-bp step) and counted how many 3’-ends fall into each window. The window with the highest number of 3’-ends was defined as the major CS, and cleavage strength was calculated by dividing this number with the number of 3’-ends that fall into the whole pA.

#### High-quality convergent gene pairs

Convergent gene pair means two neighboring protein-coding genes are located on the opposite strand and are transcribed towards each other. To check the 3’UTR overlapping pattern between convergent gene pairs, we discarded those pairs: 1) either one doesn’t have an assigned pA downstream of stop codon; 2) either one is a dubious or an overlapping gene. Finally, we got 1,396 high-quality convergent gene pairs.

#### Visualization of bidirectional termination pattern

Protein-coding genes were first separated into convergent and tandem genes. Each group was further separated into single-UTR genes (only 1 pA) and multi-UTR genes (>1 pAs) (pAs that are within CDR and generate truncated transcripts were not used in this analysis). For single-UTR genes, they were aligned by the position of CSs (coordinate 0), while multi-UTR genes were aligned by the position of distal (most distant to stop codon) CSs. Then we counted how many 3’-ends (position of last nucleotide) of AS reads fall into each position of a 1000 bp window centered at coordinate 0 (left side: upstream; right side: downstream). These count matrixes were ranked by either 3’UTR length (for single-UTR genes) or distance between proximal (closest to stop codon) and distal CSs (for multi-UTR genes). Then they were z-scale transformed and visualized by heatmaps.

#### Visualization of nucleosome coverage pattern

For convergent genes, this analysis is similar to the above-mentioned visualization of bidirectional termination pattern, except for nucleosome coverage data were used instead of 3’-ends of AS reads. For tandem genes, the count matrixes were ranked by the distance between CSs (or distal CSs for multi-UTR genes) and downstream TSSs, instead of by 3’UTR length or the distance between proximal and distal CSs. Coordinates of TSSs were downloaded from published dataset^48^.

#### Expression correlation within bidirectional termination zones

For pairs of sense and AS pAs in bidirectional termination zones, they were grouped by the relative distance between sense and AS CSs. For example, coordinate 0 means sense and AS CSs align at the same location, and coordinate -30 mean that AS CS is located at 30 bp upstream of sense CS. For each distance group, a Spearman’s rho was calculated between expression levels of sense and AS pAs. To boost confidence of correlation analysis, we used a 15-bp sliding window (3-bp step) centered at each position (for example, correlation analysis for coordinate 0 includes all pA pairs of which the AS CS falls into [-7, 7] bp region around sense CS).

## Data availability

3’-seq data have been deposited in Gene Expression Omnibus and will be published with a separate manuscript. Tif-seq data (SAM files) was downloaded from website: http://steinmetzlab.embl.de/TIFSeq/. Other datasets were downloaded from NCBI (nanopore mRNA-direct sequencing data: PRJNA398797^19^ and PRJNA408327^20^; strand-specific RNA-seq: PRJNA245106^49^ and GSE69384^50^; 3’READs data: GSE151196^38^ and GSE95139^16^; Pab1 CLIP-seq: GSE46742^36^; nucleosome coverage data: GSE97290^46^).

## Acknowledgements

We thank Christine Mayr for sharing the unpublished 3’-seq data. We thank Christine Mayr and Mayr lab for their helpful discussion during this project. We thank Christine Mayr, Mervin M. Fansler, and Sibylle Mitschka for their critical comments on the manuscript.

## Authors’ Contributions

G. Z. conceived the project, analyzed data, and wrote the manuscript. G.Z and B.K. preprocessed the data. B. K. carried out the 3’-seq experiments and gathered all public datasets. Both authors edited the manuscript.

## Declaration of interests

The authors have no competing interests.

## Notes

### Competing Interest Statement

The authors have declared no competing interest.

## Reference

1. David, L. et al. A high-resolution map of transcription in the yeast genome. Proc. Natl. Acad. Sci. U.S.A. 103, 5320–5325 (2006).

2. Pelechano, V., Wei, W. & Steinmetz, L. M. Extensive transcriptional heterogeneity revealed by isoform profiling. Nature 497, 127–131 (2013).

3. de Boer, C. G. et al. A unified model for yeast transcript definition. Genome Res. 24, 154–166 (2014).

4. Xu, Z. et al. Bidirectional promoters generate pervasive transcription in yeast. Nature 457, 1033–1037 (2009).

5. Neil, H. et al. Widespread bidirectional promoters are the major source of cryptic transcripts in yeast. Nature 457, 1038–1042 (2009).

6. Schulz, D. et al. Transcriptome surveillance by selective termination of noncoding RNA synthesis. Cell 155, 1075–1087 (2013).

7. Kuehner, J. N., Pearson, E. L. & Moore, C. Unravelling the means to an end: RNA polymerase II transcription termination. Nat Rev Mol Cell Biol 12, 283–294 (2011).

8. Irniger, S., Egli, C. M. & Braus, G. H. Different classes of polyadenylation sites in the yeast Saccharomyces cerevisiae. MOL. CELL. BIOL. 11, 3060–3069 (1991).

9. Egli, C. Sequence requirements of the bidirectional yeast TRP4 mRNA 3’-end formation signal. Nucleic Acids Research 25, 417–422 (1997).

10. Nagalakshmi, U. et al. The transcriptional landscape of the yeast genome defined by RNA sequencing. Science 320, 1344–1349 (2008).

11. Wang, L. et al. 3′ untranslated regions mediate transcriptional interference between convergent genes both locally and ectopically in Saccharomyces cerevisiae. PLoS Genet 10, e1004021 (2014).

12. Uwimana, N., Collin, P., Jeronimo, C., Haibe-Kains, B. & Robert, F. Bidirectional terminators in Saccharomyces cerevisiae prevent cryptic transcription from invading neighboring genes. Nucleic Acids Research 45, 6417–6426 (2017).

13. Mayr, C. Evolution and biological roles of alternative 3′UTRs. Trends in Cell Biology 26, 227–237 (2016).

14. Elkon, R., Ugalde, A. P. & Agami, R. Alternative cleavage and polyadenylation: extent, regulation and function. Nature Reviews Genetics 14, 496–506 (2013).

15. Ozsolak, F. et al. Comprehensive polyadenylation site maps in yeast and human reveal pervasive alternative polyadenylation. Cell 143, 1018–1029 (2010).

16. Liu, X. et al. Comparative analysis of alternative polyadenylation in S. cerevisiae and S. pombe. Genome Res. 27, 1685–1695 (2017).

17. Brooks, A. N. et al. Transcriptional neighborhoods regulate transcript isoform lengths and expression levels. Science 375, 1000–1005 (2022).

18. Lianoglou, S., Garg, V., Yang, J. L., Leslie, C. S. & Mayr, C. Ubiquitously transcribed genes use alternative polyadenylation to achieve tissue-specific expression. Genes & Development 27, 2380–2396 (2013).

19. Jenjaroenpun, P. et al. Complete genomic and transcriptional landscape analysis using third-generation sequencing: a case study of Saccharomyces cerevisiae CEN.PK113-7D. Nucleic Acids Research 46, e38– e38 (2018).

20. Garalde, D. R. et al. Highly parallel direct RNA sequencing on an array of nanopores. Nature Methods 15, 201–206 (2018).

21. Schlackow, M. et al. Genome-wide analysis of poly(A) site selection in Schizosaccharomyces pombe. RNA 19, 1617–1631 (2013).

22. Mangone, M. et al. The landscape of C. elegans 3′UTRs. Science 329, 432–435 (2010).

23. Till, P., Mach, R. L. & Mach-Aigner, A. R. A current view on long noncoding RNAs in yeast and filamentous fungi. Appl Microbiol Biotechnol 102, 7319–7331 (2018).

24. Long Non Coding RNA Biology. vol. 1008 (Springer Singapore, 2017).

25. Balarezo-Cisneros, L. N. et al. Functional and transcriptional profiling of non-coding RNAs in yeast reveal context-dependent phenotypes and in trans effects on the protein regulatory network. PLoS Genet 17, e1008761 (2021).

26. Statello, L., Guo, C.-J., Chen, L.-L. & Huarte, M. Gene regulation by long non-coding RNAs and its biological functions. Nat Rev Mol Cell Biol 22, 96–118 (2021).

27. Mercer, T. R., Dinger, M. E. & Mattick, J. S. Long non-coding RNAs: insights into functions. Nat Rev Genet 10, 155–159 (2009).

28. van Dijk, E. L. et al. XUTs are a class of Xrn1-sensitive antisense regulatory non-coding RNA in yeast. Nature 475, 114–117 (2011).

29. Martens, J. A., Laprade, L. & Winston, F. Intergenic transcription is required to repress the Saccharomyces cerevisiae SER3 gene. Nature 429, 571–574 (2004).

30. van Werven, F. J. et al. Transcription of two long noncoding RNAs mediates mating-type control of gametogenesis in budding yeast. Cell 150, 1170–1181 (2012).

31. Zhao, J., Hyman, L. & Moore, C. Formation of mRNA 3′ ends in eukaryotes: mechanism, regulation, and interrelationships with other steps in mRNA synthesis. Microbiol. Mol. Biol. Rev. 63, 405–445 (1999).

32. Irniger, S. & Braus, G. H. Saturation mutagenesis of a polyadenylation signal reveals a hexanucleotide element essential for mRNA 3’ end formation in Saccharomyces cerevisiae. Proc. Natl. Acad. Sci. U.S.A. 91, 257–261 (1994).

33. Guo, Z. & Sherman, F. 3’-end-forming signals of yeast mRNA. Mol. Cell. Biol. 15, 5983–5990 (1995).

34. Shalem, O. et al. Systematic dissection of the sequence determinants of gene 3’ end mediated expression control. PLOS Genetics 11, e1005147 (2015).

35. Savinov, A., Brandsen, B. M., Angell, B. E., Cuperus, J. T. & Fields, S. Effects of sequence motifs in the yeast 3′ untranslated region determined from massively parallel assays of random sequences. Genome Biol 22, 293 (2021).

36. Tuck, A. C. & Tollervey, D. A transcriptome-wide atlas of RNP composition reveals diverse classes of mRNAs and lncRNAs. Cell 154, 996–1009 (2013).

37. Levin, J. Z. et al. Comprehensive comparative analysis of strand-specific RNA sequencing methods. Nature Methods 7, 709–715 (2010).

38. Geisberg, J. V., Moqtaderi, Z. & Struhl, K. The transcriptional elongation rate regulates alternative polyadenylation in yeast. eLife 9, e59810 (2020).

39. Amrani, N., Minet, M., Le Gouar, M., Lacroute, F. & Wyers, F. Yeast Pab1 interacts with Rna15 and participates in the control of the poly(A) tail length in vitro. Mol Cell Biol 17, 3694–3701 (1997).

40. Kessler, S. H. & Sachs, A. B. RNA recognition motif 2 of yeast Pab1p is required for its functional interaction with eukaryotic translation initiation factor 4G. Mol Cell Biol 18, 51–57 (1998).

41. Proudfoot, N. J. Ending the message: poly(A) signals then and now. Genes Dev. 25, 1770–1782 (2011).

42. .Mavrich, T. N. et al. A barrier nucleosome model for statistical positioning of nucleosomes throughout the yeast genome. Genome Res. 18, 1073–1083 (2008).

43. Shivaswamy, S. et al. Dynamic remodeling of individual nucleosomes across a eukaryotic genome in response to transcriptional perturbation. PLoS Biol 6, e65 (2008).

44. Brogaard, K., Xi, L., Wang, J.-P. & Widom, J. A map of nucleosome positions in yeast at base-pair resolution. Nature 486, 496–501 (2012).

45. Fan, X. et al. Nucleosome depletion at yeast terminators is not intrinsic and can occur by a transcriptional mechanism linked to 3’-end formation. Proc. Natl. Acad. Sci. U.S.A. 107, 17945–17950 (2010).

46. Chereji, R. V., Ramachandran, S., Bryson, T. D. & Henikoff, S. Precise genome-wide mapping of single nucleosomes and linkers in vivo. Genome Biol 19, 19 (2018).

47. Chereji, R. V., Bryson, T. D. & Henikoff, S. Quantitative MNase-seq accurately maps nucleosome occupancy levels. Genome Biol 20, 198 (2019).

48. Park, D., Morris, A. R., Battenhouse, A. & Iyer, V. R. Simultaneous mapping of transcript ends at single-nucleotide resolution and identification of widespread promoter-associated non-coding RNA governed by TATA elements. Nucleic Acids Research 42, 3736–3749 (2014).

49. Smith, J. E. et al. Translation of small open reading frames within unannotated RNA transcripts in Saccharomyces cerevisiae. Cell Reports 7, 1858–1866 (2014).

50. Wery, M. et al. Nonsense-mediated decay restricts lncRNA levels in yeast unless blocked by double-stranded RNA structure. Molecular Cell 61, 379–392 (2016).

51. Lengronne, A. et al. Cohesin relocation from sites of chromosomal loading to places of convergent transcription. Nature 430, 573–578 (2004).

52. Paldi, F. et al. Convergent genes shape budding yeast pericentromeres. Nature 582, 119–123 (2020).

53. Nishiyama, T. Cohesion and cohesin-dependent chromatin organization. Current Opinion in Cell Biology 58, 8–14 (2019).

54. Hou, Y. & Lin, S. Distinct gene number-genome size relationships for eukaryotes and non-eukaryotes: gene content estimation for dinoflagellate genomes. PLoS ONE 4, e6978 (2009).

55. Ju, X., Li, D. & Liu, S. Full-length RNA profiling reveals pervasive bidirectional transcription terminators in bacteria. Nat Microbiol 4, 1907–1918 (2019).

56. Bolger, A. M., Lohse, M. & Usadel, B. Trimmomatic: a flexible trimmer for Illumina sequence data. Bioinformatics 30, 2114–2120 (2014).

57. Martin, M. Cutadapt removes adapter sequences from high-throughput sequencing reads. EMBnet.journal 17, 10–12 (2011).

58. Kim, D., Paggi, J. M., Park, C., Bennett, C. & Salzberg, S. L. Graph-based genome alignment and genotyping with HISAT2 and HISAT-genotype. Nat Biotechnol 37, 907–915 (2019).

59. Derti, A. et al. A quantitative atlas of polyadenylation in five mammals. Genome Res. 22, 1173–1183 (2012).

60. Li, H. Minimap2: pairwise alignment for nucleotide sequences. Bioinformatics 34, 3094–3100 (2018).

61. Quinlan, A. R. & Hall, I. M. BEDTools: a flexible suite of utilities for comparing genomic features. Bioinformatics 26, 841–842 (2010).

62. Liao, Y., Smyth, G. K. & Shi, W. featureCounts: an efficient general purpose program for assigning sequence reads to genomic features. Bioinformatics 30, 923–930 (2014).

63. Mainul, H., Li, W. & Tian, B. Accurate mapping of cleavage and polyadenylation sites by 3′ region extraction and deep sequencing. In: Rorbach, J., Bobrowicz, A. (eds) Polyadenylation. Methods in Molecular Biology 1125, (2014).

