## Supplemental figures for "Transcription is ubiquitously terminated in thousands of bidirectional termination zones in yeast"

#### Supplemental figure legends

Supp. Figure 1: Additional information on pA annotation of protein-coding genes. (a) one pA located in coding region of BRR6 is in fact a termination site of the upstream gene PDE1. (b) Two pAs downstream of YBR241C in fact belong to ncRNA XUT_2R-296. (c) pAs belong to NCS6 and YPT32 are influenced by polycistronic reads, making proper downstream analysis (e.g., differential expression analysis) quite tricky. (d) Venn diagram shows pA assignment results are highly consistent among three long-read sequencing data (Nanopore 1: Garalde, Nature Methods, 2018; Nanopore 2: Jenjaroenpun, NAR, 2018; Tif-seq: Pelechano, Nature, 2013). (e) Number of pAs fall into 3'UTR or coding region (CDR), and number of pAs influenced by polycistronic reads. (f) 3'UTR length distribution in yeast (only 3'UTRs with length <800 bp were shown in the histogram).

Supp. Figure 2: Additional information on pA annotation of non-coding genes. (a-f) APA events of well-known ncRNAs (a: SRGs; b: IRT1), XUTs (c; XUT_2F-93; d: XUT_3R-67), and CUTs (e: CUT040; f: UT492). (g) Venn diagram shows genomic overlapping of three classes of ncRNAs in our dataset. (h) Summary of pA annotation results for three classes of ncRNAs (Multi-pA: genes with >1 pAs). (i) Expression levels for four classes of transcripts (RPM: reads per million). Due to frequent overlaps among SUTs/XUTs/CUTs (panel g), we define 567 ncRNAs that overlap with SUTs but no overlap with CUTs as SUTs, 427 ncRNAs that overlap with XUTs but overlap with neither SUTs nor CUTs as XUTs, 283 ncRNAs that overlap with CUTs but no overlap with SUTs as CUTs.

Supp. Figure 3: (a) pAs with UAUAUA motifs (located in a -75 to -25 bp region upstream of cleavage site) have higher expression levels compared to those without UAUAUA motifs. (b) Examples to show pAs with a strong CS (top panel, most transcripts are cut at the same position, high cleavage strength) and a weak CS (bottom panel, transcripts are cut at multiple sites, weak cleavage strength). (c) UAUAUA motif's position relative to CS can influence cleavage strength. (d) AAUAAA motif distributions among different pA categories (single/distal/proximal) and among different classes of transcripts (mRNAs/SUTs/XUTs/CUTs). (e) Ratios of transcripts with AAUAAA motif (located in [-30, -5] region) and UAUAUA motif (located in [-75, -25] region).

Supp. Figure 4: (a) 3'UTR overlapping of convergent genes MRPL32 and YCP4. (b) 3'UTR overlapping of convergent genes SUB2 and RPS16B. (c-h) 3'-end distribution of AS reads centered at CSs of convergent genes with bidirectional terminators using three other datasets (c,d: 3'-READs data (Geisberg, Elife, 2020); e,f: Tif-seq data (Pelechano, Nature, 2013); g,h: Pab1 CLIP-seq data (Tuck, Cell, 2013)). (c, e, g) Single-UTR genes. (d, f, h) Multi-UTR genes.

Supp. Figure 5: (a) One example shows two convergently transcribed coding genes (SGE1 and ARR1) don't share their 3'UTRs, yet they share their 3'UTRs with termination sites of two AS ncRNAs. (b, c) 3'-end distribution of A-S nanopore reads centered at CSs of convergent genes that share termination sites with AS ncRNAs (b: single-UTR genes; c: multi-UTR genes). Black dots in (c) represent positions of proximal CSs.

Supp. Figure 6: 3'-end distribution of AS reads centered at CSs of unidirectional convergent genes. (a, b) AS reads from nanopore data (Jenjaroenpun, NAR, 2018). (c, d) AS reads from 3'-READs data (Geisberg, Elife, 2020). (e, f) AS reads from Tif-seq (Pelechano, Nature, 2013). (g, h) AS reads from Pab1 CLIP-seq data (Tuck, Cell, 2013). (a, c, e, g) Unidirectional single-UTR genes. (b, d, f, h) Unidirectional multi-UTR genes.

Supp. Figure 7: (a, b) two examples show protein-coding genes (a: SVF1, b: NOP10) share their 3'UTR with termination sites of AS ncRNAs. (c-h) 3'-end distribution of AS reads centered at CSs of tandem genes, whose 3UTRs overlap with termination sites of AS ncRNAs, using three other datasets. (c, d) 3'-READs data (Geisberg, Elife, 2020). (e, f) Tif-seq data (Pelechano, Nature, 2013). (g, h) Pab1 CLIP-seq (Tuck, Cell, 2013). (c, e, g) Single-UTR genes. (d, f, h) Multi-UTR genes.

Supp. Figure 8: 3'-end distribution of AS nanopore mRNA-direct sequencing reads centered at CSs of tandem genes that don't share termination sites with AS ncRNAs. (a) Single-UTR genes. (b) Multi-UTR genes. Black dots represent positions of proximal CSs.

Supp. Figure 9: (a) 3'-ends of sense and AS nanopore mRNA-direct sequencing reads are symmetrically distributed on both sides of UAUAUA motifs in bidirectional termination zones of tandem genes. (b) Mono-nucleotide compositions on both sides of UAUAUA motifs in bidirectional termination zones can be reverse-complemented.

Supp. Figure 10: (a, c) Heatmaps show nucleosome occupancy in a 1000 bp window centered at CSs of convergent (a) and tandem (c) single-UTR genes with unidirectional terminators. (b, d) Heatmaps show nucleosome occupancy in a 1000 bp window centered at distal CSs of convergent (b) and tandem (d) multi-UTR genes with unidirectional terminators. Black dots in b & d represent positions of proximal CSs. Green dots in c & d represent positions of downstream TSSs. (e) For tandem genes that have bidirectional terminators (ncRNA olp.), their CSs tend to be located much further away from downstream TSSs than those of tandem genes with unidirectional terminators (no olp.).

Supp. Figure 11: (a, b) No expression correlation at gene level between convergent protein-coding genes using either 3'-READs data (a) or strand-specific RNA-seq data (b). (c) One example shows expression of sense pAs seems to have a negative relationship with expression of AS pAs when their CSs are close to each other. (d) For the 1,495 AS CSs that fall into [-60, -20] region of sense CPEs without UAUAUA motifs, there is an enrichment of UAUAUA motifs downstream of their CSs (UAUAUA motifs also include two key variants: UACAUA and UAUGUA). In (a) and (b), R values were calculated using Spearman's method.

Zhen & Buki, 2022
Supplemental Figure 1

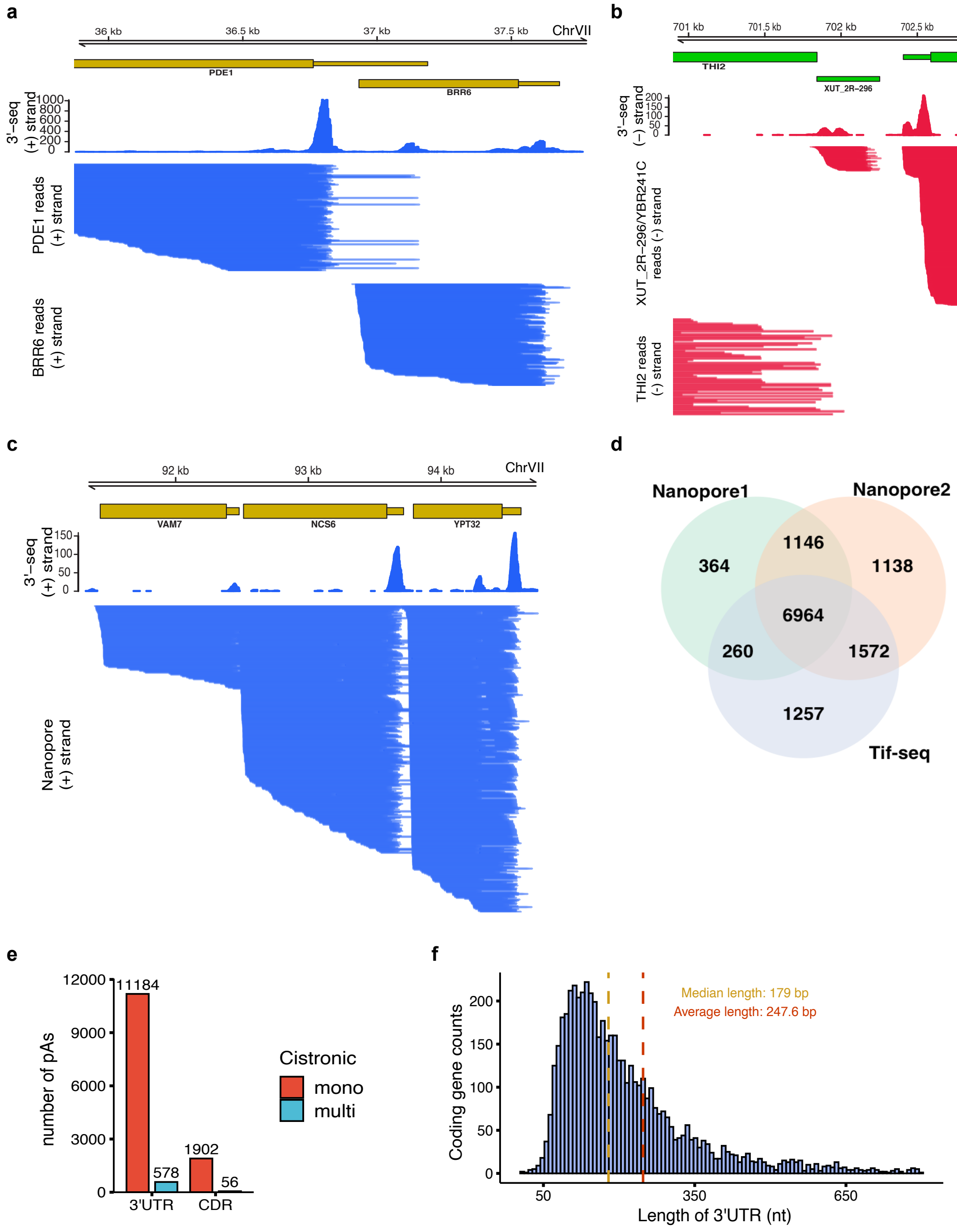

Zhen & Buki, 2022
Supplemental Figure 2

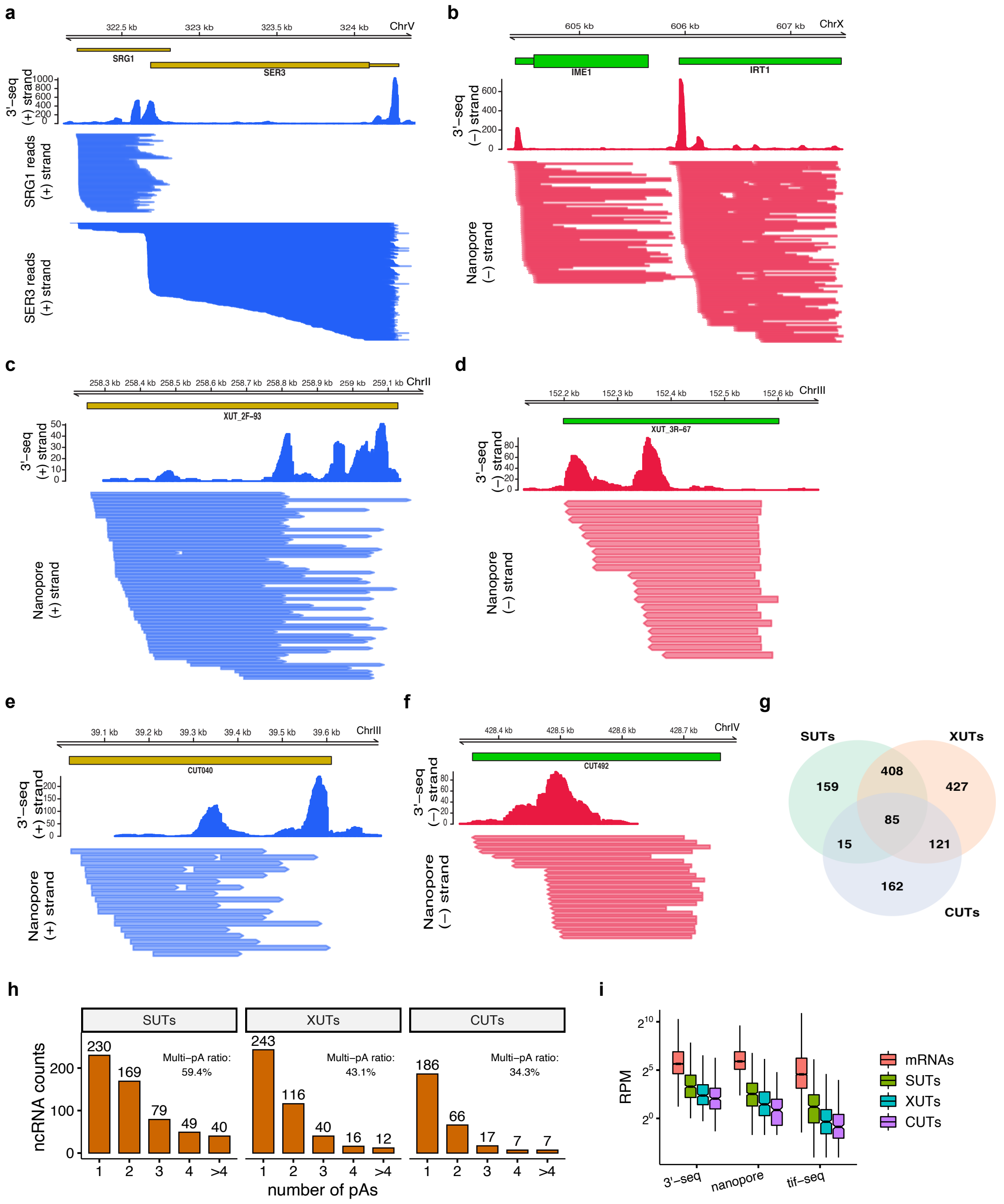

Zhen & Buki, 2022
Supplemental Figure 3

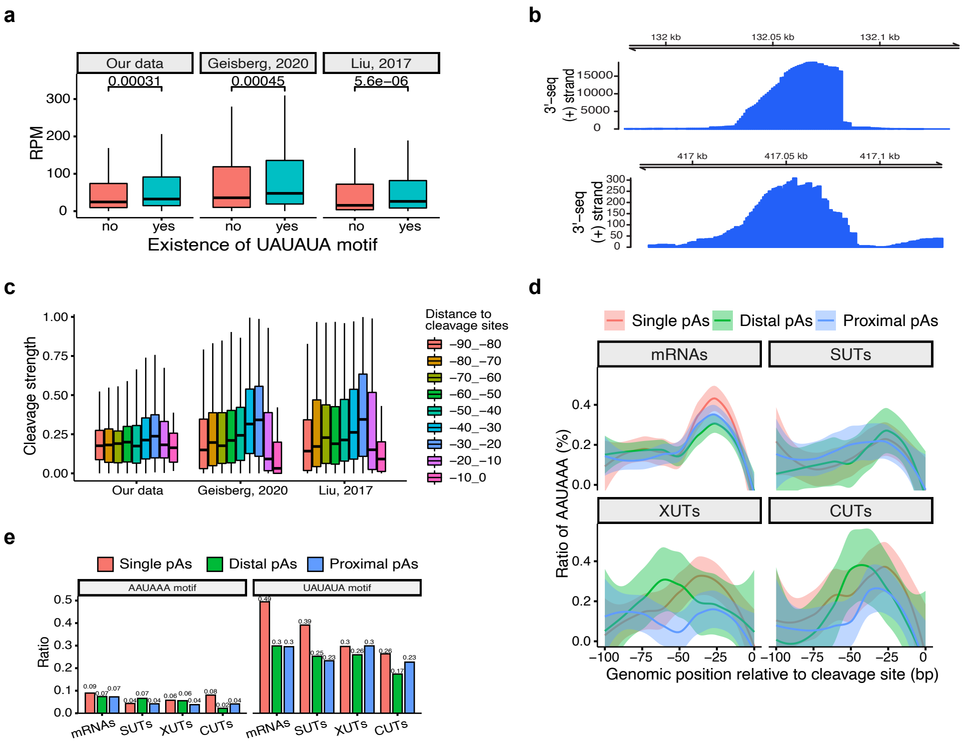

### Zhen & Buki, 2022
### Supplemental Figure 4

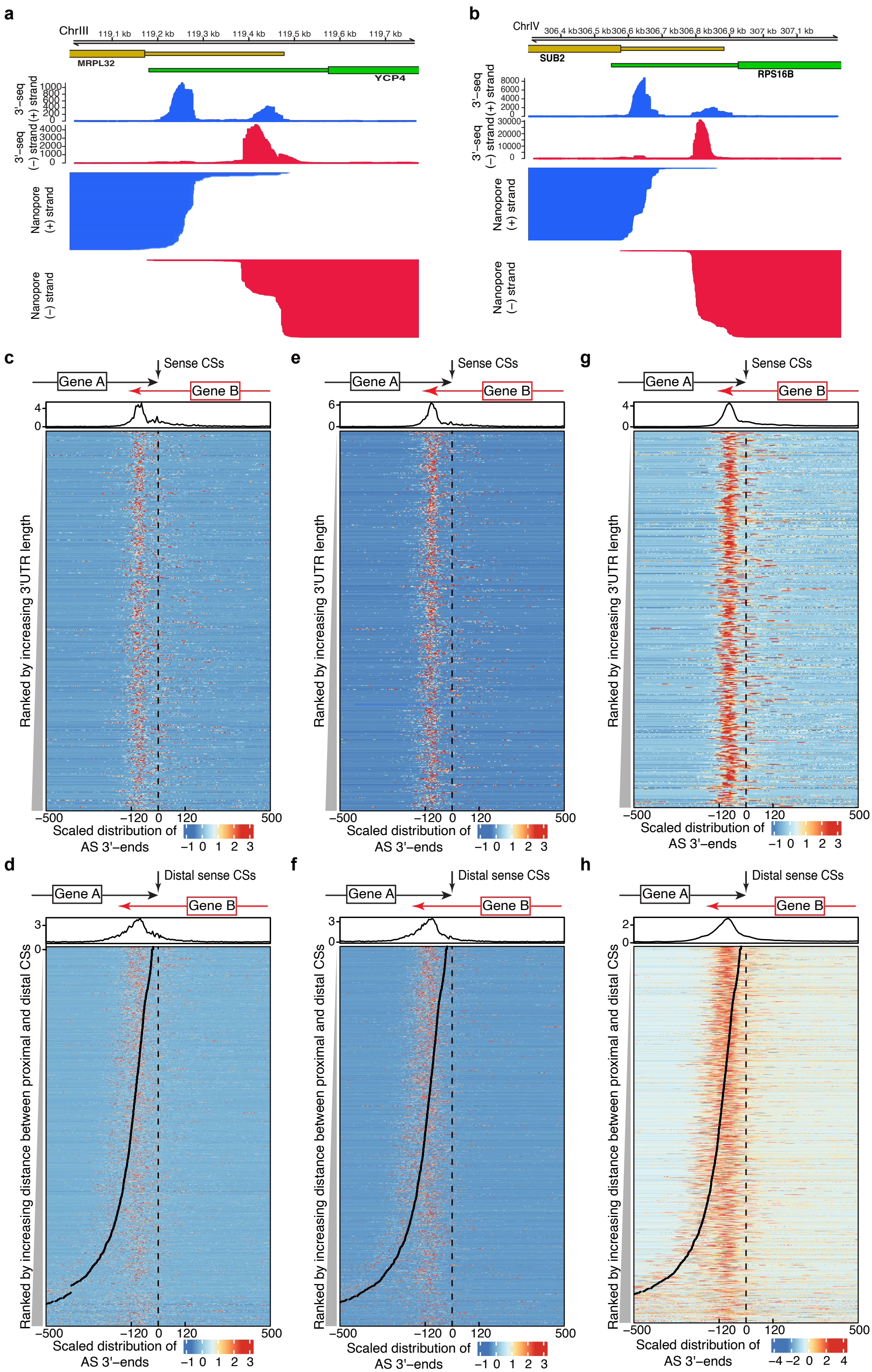

Zhen & Buki, 2022
Supplemental Figure 5

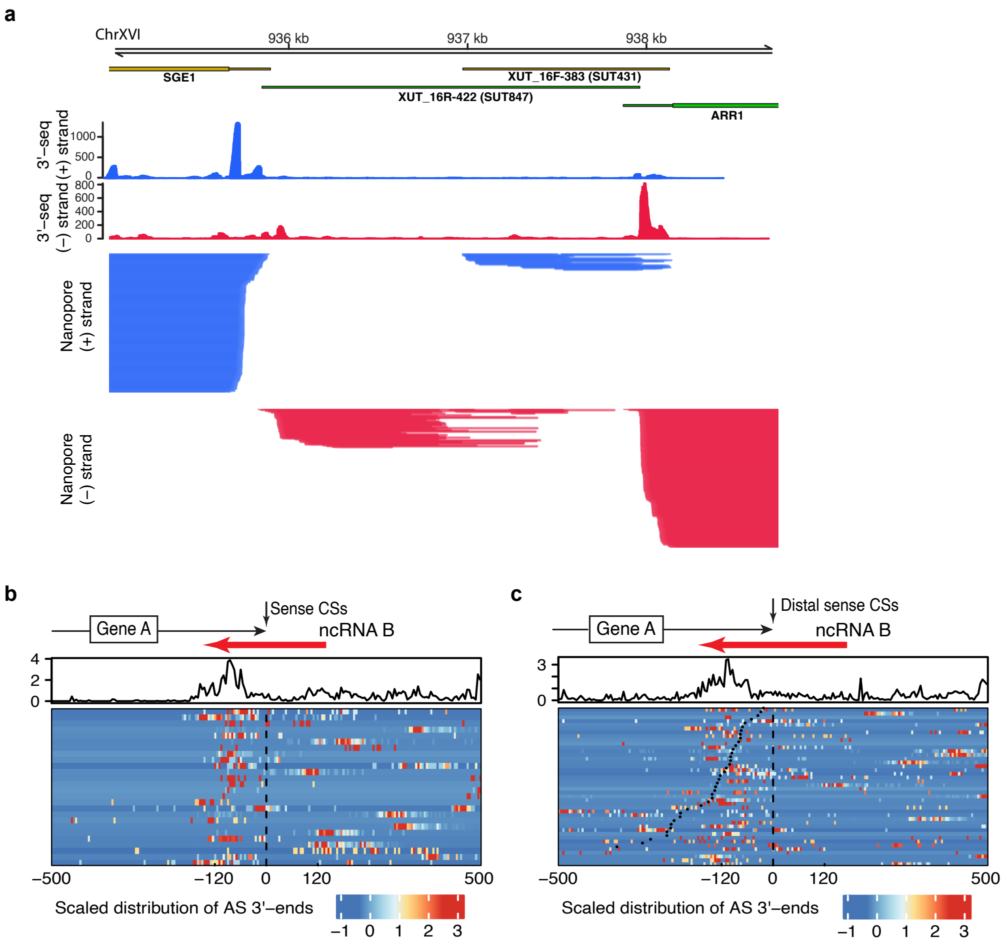

### Zhen & Buki, 2022
### Supplemental Figure 6

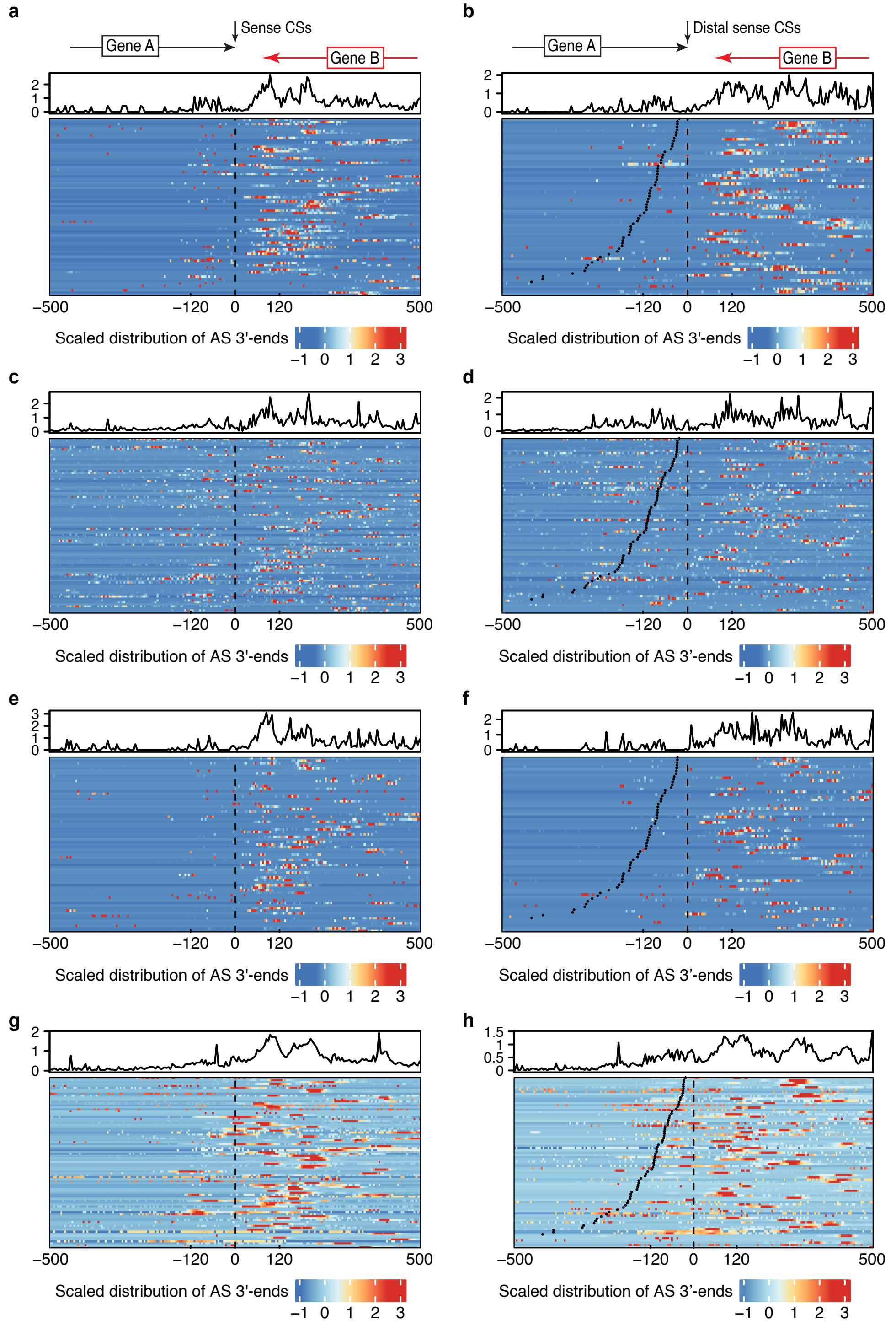

### Zhen & Buki, 2022
### Supplemental Figure 7

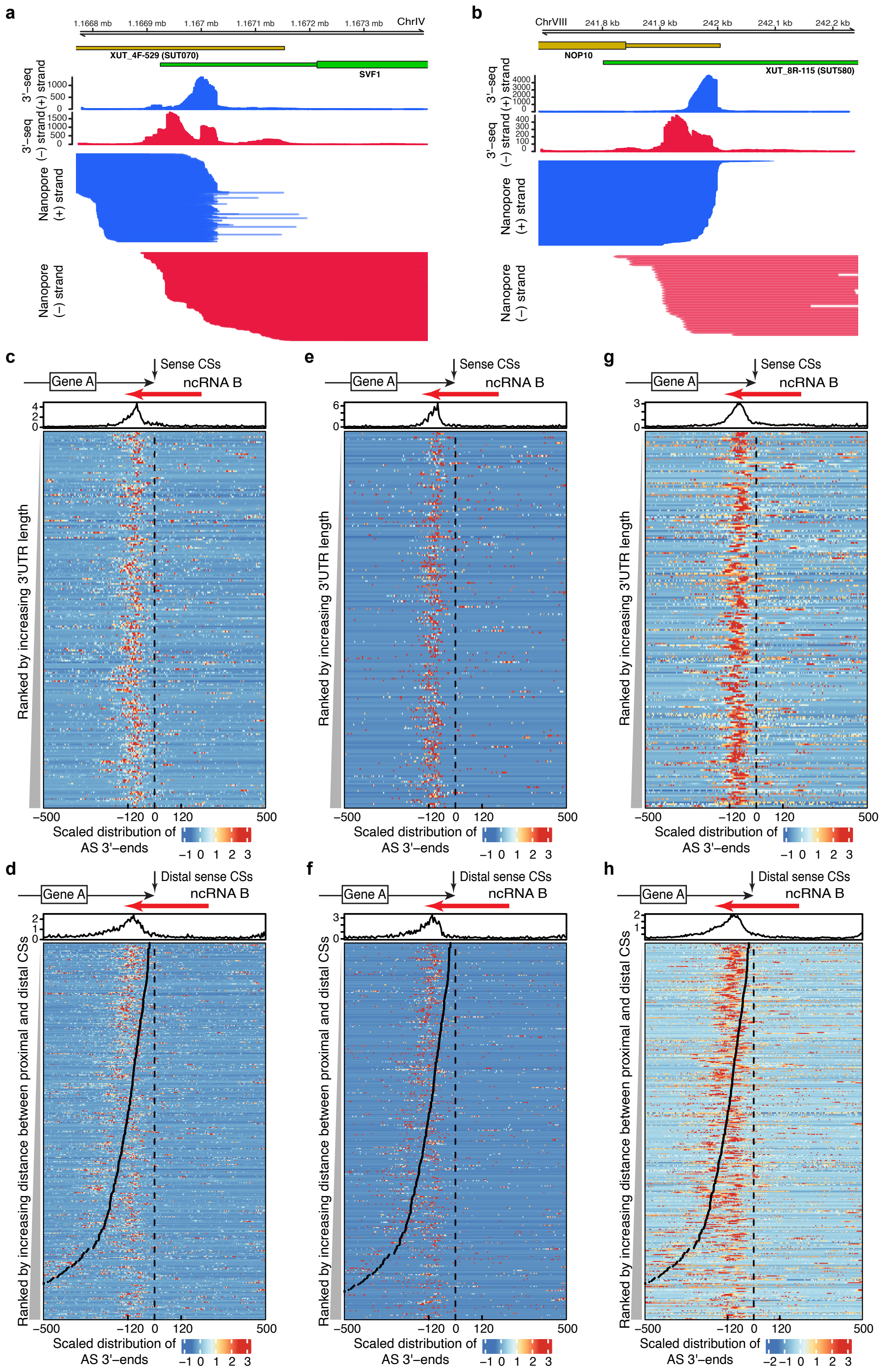

Zhen & Buki, 2022
Supplemental Figure 8

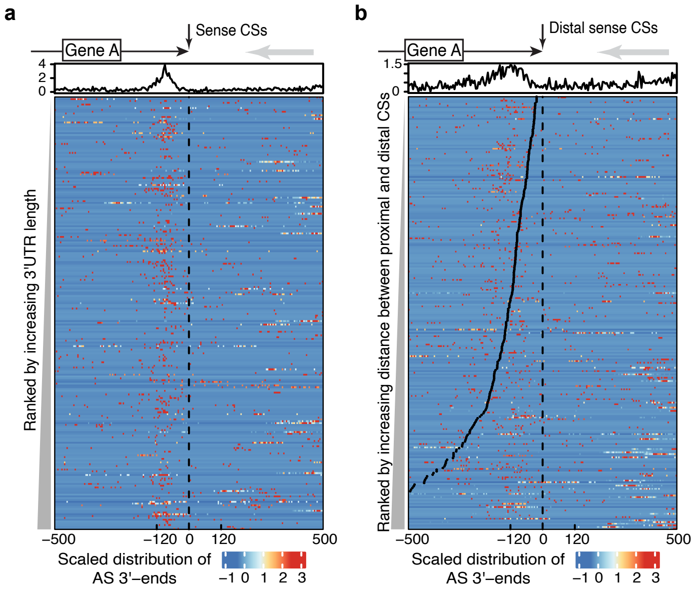

### Zhen & Buki, 2022
### Supplemental Figure 9

**a**

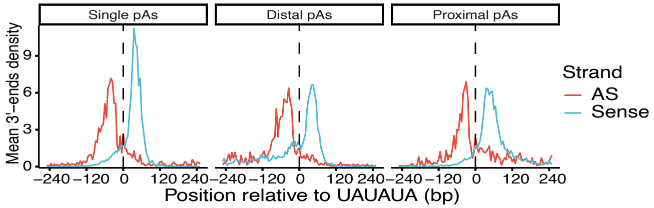


**b**

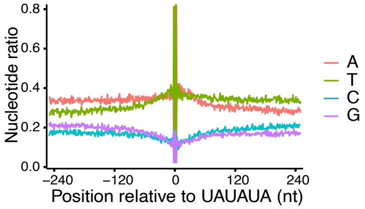

### Zhen & Buki, 2022
### Supplemental Figure 10

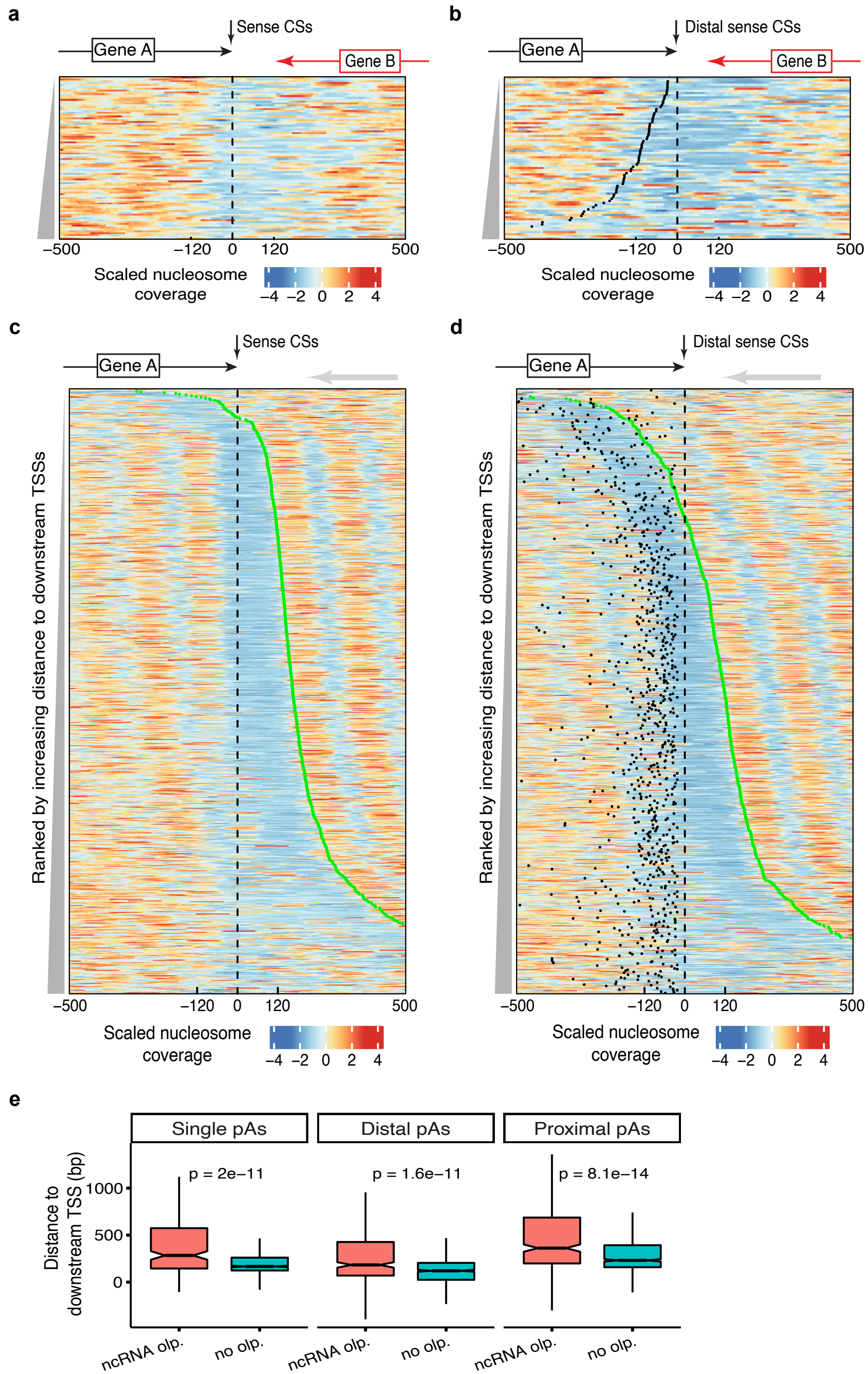

Zhen & Buki, 2022
Supplemental Figure 11

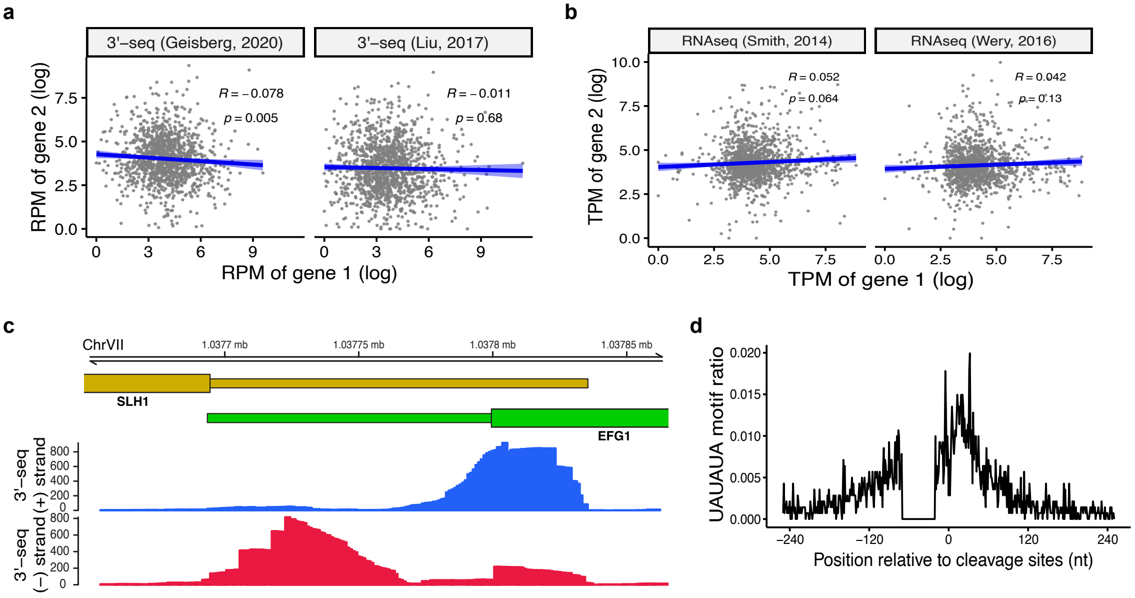
